# The 3D affinities of the OT-I TCR to foreign and self-antigens predict their 2D affinities and reveal imperfect antigen discrimination

**DOI:** 10.1101/2025.01.12.632665

**Authors:** Anna Huhn, Mikhail A. Kutuzov, Keir Maclean, Lion F. K. Uhl, Jagdish M. Mahale, Audrey Gerard, P. Anton van der Merwe, Omer Dushek

## Abstract

T cells use the T cell antigen receptor (TCR) to discriminate between higher-affinity foreign and lower-affinity self peptide-MHC (pMHC) antigens. The OT-I mouse TCR is widely used to study antigen discrimination utilising many pMHCs, including foreign and self antigens. Previous studies suggested that OT-I T cells achieve near-perfect discrimination between higher and lower affinity antigens. Moreover, these 3D affinities measured in solution did not correlate with the 2D membrane affinities, suggesting a complex relationship between 3D and 2D affinities. In contrast, other TCRs have shown imperfect antigen discrimination and strong correlations between 3D and 2D affinities. To resolve these discrepancies, we extended a protocol for measuring ultra-low TCR/pMHC affinities to accurately determine the 3D affinities of the OT-I TCR binding 19 pMHC complexes. These revised 3D affinities now strongly correlate with the 2D affinities, and accurately predict functional responses. Importantly, we now find that the OT-I TCR exhibits enhanced yet imperfect antigen discrimination, similar to other TCRs, allowing it to detect abnormally high levels of low-affinity self-antigens. Finally, we show that discrimination is highest with low-affinity pMHC ligands, a finding predicted by the kinetic-proofreading model of antigen discrimination. This work underscores the ability of T cells to effectively gauge proxies for 3D affinity within the 2D cell-cell interface, with significant implications for the mechanisms underlying antigen discrimination.

**Lay abstract:** T cells protect the body from infection by distinguishing between foreign and self molecules. They do this using a specialized receptor called the T cell receptor (TCR), which senses differences in how strongly it binds to foreign versus self molecules. Scientists have used a mouse TCR called OT-I to understand this process, concluding that OT-I T cells exhibit near perfect discrimination between foreign and self antigens based on a sharp affinity threshold. However, conflicting results from experiments on other TCRs hinted at a more complicated picture. In this study, we used an improved method to accurately measure OT-I TCR binding affinity to 19 peptides, including foreign and self antigens. In contrast to previous results and like other TCRs, we found that OT-I T cells display enhanced but imperfect discrimination, enabling them to be activated by high levels of self antigen. The new data also revealed that the ability of T cells to discriminate antigens is particularly high at low affinities. These findings reconcile apparent discrepancies between the OT-I TCR and other TCRs, and have implications for understanding diseases where T cells often respond to self antigen, such as autoimmunity and cancer.

**Graphical abstract:** 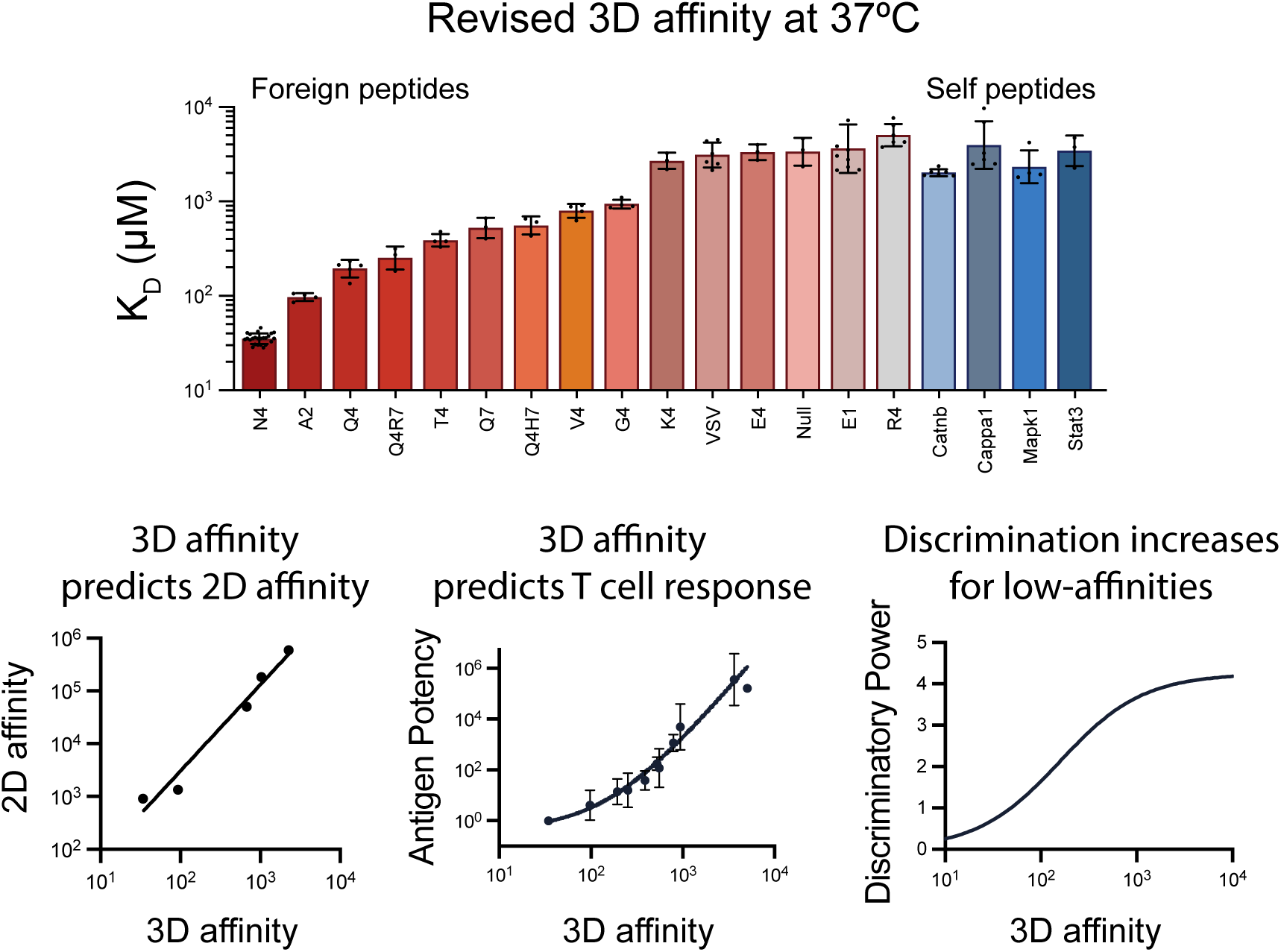

## Introduction

T cells orchestrate adaptive immune responses by recognizing infected or cancerous cells whilst ignoring normal cells. This ability relies on their T cell receptors (TCRs) being able to discriminate between higher-affinity foreign and lower-affinity self peptide antigens presented on major-histocompatibility-complexes (pMHCs) on antigen presenting cell (APC) surfaces. This process was first explored using mouse T cell hybridomas and transgenic mice expressing TCRs of defined specificity (1–7). The OT-I TCR transgenic mouse has been among the most widely used *in vivo* systems for studying T cell responses, including central and peripheral tolerance, infection, cancer, vaccination, autoimmunity, and transplantation (8–18). The OT-I TCR, which recognises the ovalbumin-derived peptide SIINFEKL (N4) presented by the MHC class I protein H-2K*^b^*, has also been used to investigate the molecular and cellular mechanisms underlying T cell antigen recognition (19–25).

During the past three decades, more than 20 additional peptides have been used to investigate how varying the peptide affects antigen recognition by OT-I T cells. Early studies suggested that the OT-I TCR exhibits near-perfect antigen discrimination (1–3, 5, 19). For example, while the OT-I TCR was reported to bind the E1 peptide with only a 3-fold lower affinity than the N4 peptide, it required a 100,000-fold higher concentration to be activated (3). This striking observation spurred extensive theoretical and experimental efforts to uncover the mechanism(s) enabling such exceptional discrimination (19, 21, 26–36). One prominent hypothesis was that solution or 3D affinities, commonly measured using surface plasmon resonance (SPR) and soluble forms of TCRs and pMHCs, may not accurately reflect the 2D affinities between membrane-attached TCRs and pMHCs, because the latter interactions are exposed to force (37). Indeed, 2D affinity measurements of OT-I/pMHC interactions revealed much larger variations, with a 200-fold difference between the N4 and E1 compared with only 3-fold difference in 3D affinity (20). Thus, the OT-I TCR apparently displays a highly non-linear relationship between 3D and 2D affinities.

However, findings with the OT-I TCR have not been replicated with other TCRs. Using an optimized SPR protocol for measuring very low affinity TCR/pMHC interactions, we have shown that human TCRs display much weaker discrimination than originally reported for the OT-I TCR, in that a 3-fold lower peptide affinity requires only a 9-fold increase in peptide concentration to activate T cells (38). Furthermore, unlike the OT-I TCR, the 3D and 2D affinities of the 1E6 produced linear correlations (39), while the 3D and 2D TCR/pMHC lifetimes for the 5c.c7 TCR were comparable (40). These discrepancies between the OT-I TCR and other TCRs remain unexplained.

One advantage of the OT-I TCR is that some of the self-peptides that this TCR binds (e.g. Catnb and Cappa1) have been identified, based of their ability to positively select OT-I thymocytes (41). Given that mature OT-I T cells must ignore these self-peptides, measuring the affinities of OT-I binding these peptides would provide insights into the level of TCR discrimination required for tolerance. Unfortunately it has not been possible to measure these affinities at physiological temperatures; K_D_ estimates have only been reported at 10*^◦^*C (10).

Concentrating soluble proteins to the levels required for studying very low affinity interactions by SPR often results in formation of protein aggregates, which bind with slow kinetics (42, 43). In early SPR studies the OT-I TCR showed biphasic binding to N4 pMHC, with one component binding with unusually slow kinetics (*k*_on_ ∼ 0.03 *µ*M *^−^*^1^ s*^−^*^1^, *k*_off_ ∼ 0.02 s*^−^*^1^ for the slow phase) (3). This gave rise to the notion that the OT-I TCR interaction with N4 has unusually high affinity. However, this slow phase is also consistent with the presence of OT-I TCR aggregates. This is supported by subsequent studies that reported monophasic binding with much faster kinetics to N4 (11, 44, 45). Collectively, this suggests that inaccurate 3D affinity measurements could explain the apparent discrepancies between the OT-I TCR and other TCRs.

These discrepancies and lack of affinity data for most OT-I peptides motivated us to use our optimised SPR protocol to measure 3D affinities between the OT-I TCR and 20 commonly used peptides at 37*^◦^*C (38). We now report that the OT-I TCR binds N4 with physiological affinity and displays a much wider variations in affinity for various peptides, such as a 100-fold lower affinity for E1 relative to N4 rather than the originally reported 3-fold difference. Our revised 3D K_D_ values correlate with 2D K_D_ values, and indicate that the discriminatory power of the OT-I TCR is comparable to other TCRs, and increases for lower-affinity antigens. These findings reconcile apparent discrepancies between the OT-I TCR and other TCRs, and have important implications for understanding TCR antigen discrimination.

## Results

### Systematic measurements of OT-I TCR affinities at 37***^◦^***C

We selected a panel of 20 peptides commonly used in the literature for OT-I TCR experiments, including positively-selecting self-peptides (Table 1). Purified OT-I TCR was injected over surfaces immobilised with each pMHC at 37*^◦^*C (Fig. 1). The association and dissociation phases were too fast to allow rate constants to be estimated and with the exception of the N4 pMHC, binding did not saturate at the OT-I TCR concentration range tested. This is consistent with weak interactions. As a result, the K_D_ could not be accurately determined by conventional fitting of the 1:1 binding model where the maximal TCR binding (B_max_) is unconstrained.

**Figure 1:**
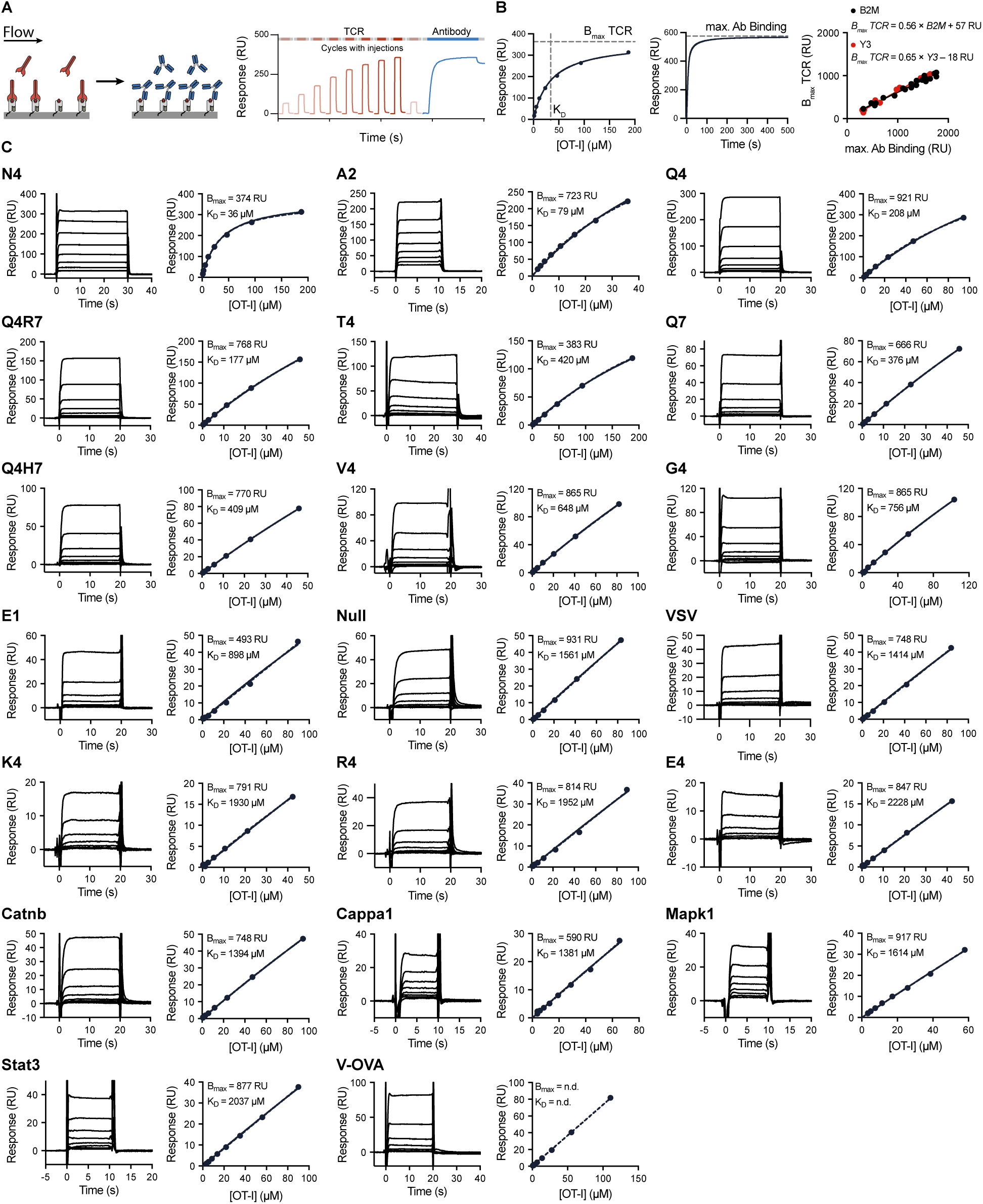
**The OT-I TCR interaction with 20 commonly used peptides exhibits fast kinetics and low-affinity at 37***^◦^***C. (A)** Schematic of SPR protocol. The TCR analyte is injected at 8 different concentrations over a surface coupled with purified pMHC followed by an injection of a pMHC conformationally sensitive antibody (Y3 or B2M). **(B)** Steady-state binding response for the higher-affinity N4 pMHC fitted with a one-site specific binding model to determine TCR B_max_ and K_D_ (left). Representative sensorgram of the B2M antibody specific for human *β*2m domain injected after the final TCR injection to obtain maximum antibody binding (centre). Empirical standard curve relating maximum antibody binding (x-axis) to fitted TCR B_max_ obtained from N4 pMHC across N=23 (B2M) and N=14 (Y3) independent experiments with different levels of pMHC on the surface (right). **(C)** Representative SPR traces (left) and steady-state binding plots (right) for the indicated peptide, Steady state data was fitted with a 1:1 binding model with constrained B_max_ to obtain K_D_.

**Table 1:**
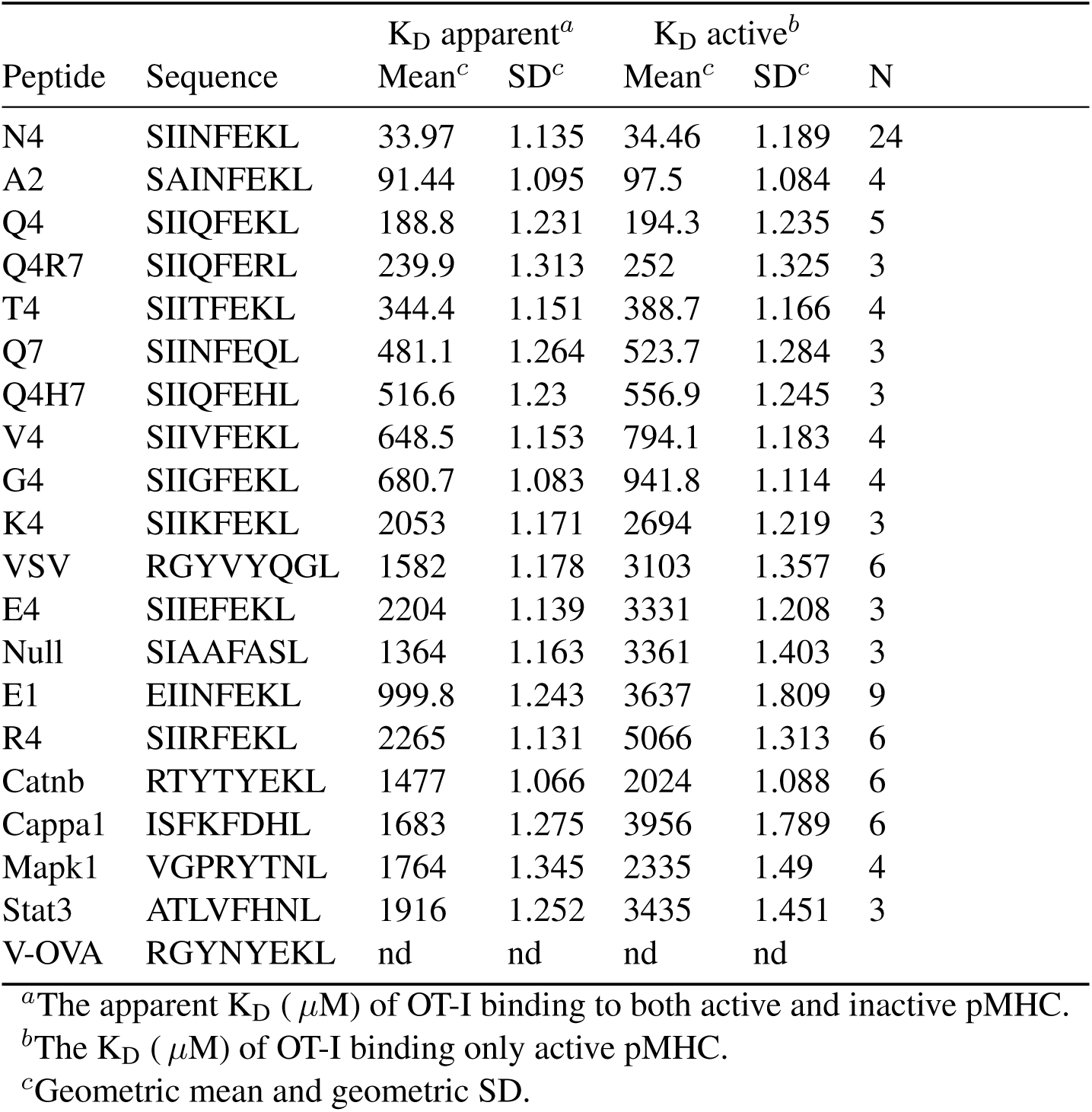
Revised K_D_ values for OT-I specific peptides at 37*^◦^*C.

To avoid concentrating the TCR, which can introduce protein aggregates, we used a previously-described SPR protocol that does not require TCR binding to approach saturation (38). We first determined the maximum TCR binding (B_max_) for surfaces immobilised with different levels of the higher-affinity N4 pMHC, where TCR binding saturates at attainable concentrations of TCR (Fig. 1A-B). We then injected the pMHC-specific B2M or Y3 antibody enabling us to produce a standard curve that relates antibody binding to the TCR B_max_ (Fig. 1B). For low-affinity TCR/pMHC interactions, where TCR binding does not saturate, measuring the B2M or Y3 antibody binding after each experimental run enabled us to use this standard curve to determine the TCR B_max_. This in turn allowed us to estimate the K_D_ by fitting the usual 1:1 binding model while constraining the value of B_max_. Using this protocol, we measured the apparent affinities for all OT-I peptides, which ranged from a K_D_ of 34 *µ*M to over 2200 *µ*M (Fig. 1C, Fig. S1, Table 1). We confirmed that pMHC produced in bacterial (*E. coli*) and mammalian (HEK293) cells produced the same affinities (Fig. S2).

Because the pMHC antibody displayed only modest binding to V-OVA [*<*30 RU versus *>*600 RU for N4 (Fig. 2A)], we were unable to determine a K_D_ for the OT-I TCR binding this peptide. This lack of binding of a conformationally sensitive antibody indicated that the V-OVA pMHC is not correctly folded, and suggested that the limited OT-I binding to V-OVA pMHC may be non-specific (Fig. 2A). To explore this, we tested OT-I binding to N4 pMHC after the latter had been denatured by exposure to low-pH glycine solution. This resulted in a 100-fold reduction in binding of the B2M antibody, confirming denaturation (Fig. 2A), yet the OT-I TCR continued to display up to 60 RU of binding to denatured N4 (compared to 400 RU to correctly-folded N4). This suggests that incorrectly folded pMHC on the sensor surface can non-specifically bind injected analytes, including the OT-I TCR. In support of this, a control protein, Ovalbumin, also showed binding to immobilised pMHCs but not to another immobilised protein, CD86 (Fig. S3). Thus, some OT-I TCR binding detected by SPR represents binding to inactive/unfolded pMHC. This non-specific binding needs to be taken into account in order to accurately measure the affinities of OT-I TCR for specific, or active, pMHCs.

**Figure 2:**
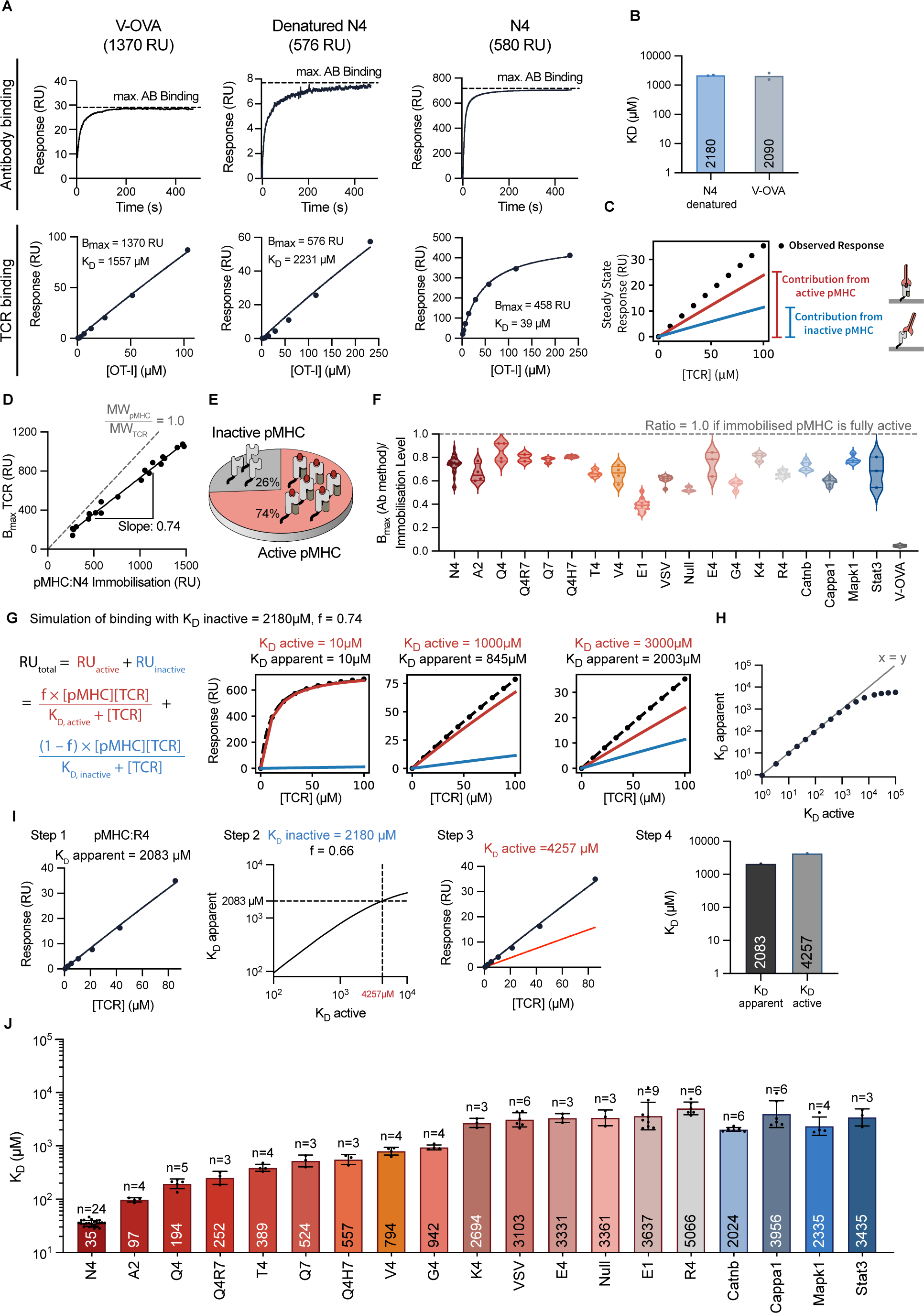
The presence of inactive pMHC can impact TCR binding and K_D_ estimates for low-affinity interactions. **(A)** SPR sensogram of B2M antibody binding (top) and OT-I TCR steady-state binding curve (bottom). The steady-state binding for denatured N4 and V-OVA were fit with a 1:1 model to estimate K_D_ with B_max_ constrained to the pMHC immobilisation level. A low pH glycine solution denatured N4. **(B)** Estimated K_D_ for the data in (A) for N=2 independent experiments. **(C)** Schematic of overall observed binding decomposed into the contribution from active and inactive pMHC. **(D)** The OT-I B_max_ over the immobilisation level of N4 produces a slope of 0.74. A slope of 1.0 is expected for a 1:1 interaction if all pMHC (49 kDa) is active and can bind the TCR (51 kDa). **(E)** Schematic showing that 74% of N4 pMHC is active and can bind the TCR. **(F)** Ratio of B_max_ (obtained from the standard curve) to pMHC immobilisation determines the fraction of active pMHC for all peptides tested. **(G)** Simulated TCR binding to surfaces with inactive and active pMHC using a fixed fraction of active pMHCs with different affinities (columns). The overall binding (black) is fit to a 1:1 model to estimate the apparent K_D_. **(H)** The fitted apparent K_D_ over the true active K_D_. **(I)** Workflow applied to convert the apparent K_D_ of OT-I/R4 interaction (2083 *µ*M) into the active K_D_ (4257 *µ*M). **(J)** Revised OT-I active K_D_ values for the indicated peptide at 37*^◦^*C (geometric mean K_D_ indicated within bar, also see Table 1 for geometric mean K_D_ +/-geometric SD.)

To estimate the OT-I affinity for inactive V-OVA and denatured N4, we fit the steady-state data with the usual 1:1 binding model but constrained the B_max_ to the total amount of pMHC immobilised (Fig. 2A, bottom). If we assume that almost all the immobilized V-OVA and denatured N4 is inactive, the immobilisation level can serve as a proxy for total available non-specific binding (B_max_) because the molecular weights of pMHC (49 kDa) and OT-I TCR (51 kDa) are nearly identical. Using this method, we found that the OT-I TCR bound denatured N4 and V-OVA with K_D_ values of 2180 and 2090 *µ*M, respectively (Fig. 2B). While this very weak binding is unlikely to affect K_D_ estimates when the fraction of inactive pMHC is very low, it would be expected to have an impact when the fraction of inactive pMHC is large (Fig. 2C).

We next estimated the fraction of inactive pMHC in the pMHC preparations. In the case of the high-affinity N4 pMHC, we can directly estimate this fraction by comparing the TCR B_max_ to the amount of immobilised N4 pMHC, which includes both active and inactive pMHC (Fig. 2D). If all pMHCs were active, B_max_ and pMHC immobilisation levels should match producing a slope of 1.0. Instead the slope of the B_max_ vs pMHC plot was 0.74, indicating that 74% of N4 is active (Fig. 2D,E). For lower affinity pMHCs the binding of conformationally sensitive antibodies was used to estimate the B_max_. The ratio of B_max_ to pMHC immobilisation indicated that the amount of active pMHC varied from 50% (E1) to 80% (Q4) (Fig. 2F).

We next modelled the effect of non-specific binding on estimates OT-I TCR K_D_ for active pMHC by extending the 1:1 binding model to include a second term to account for binding to inactive pMHC (Fig. 2G). This showed that, while having some inactive pMHCs (26%) would not distort K_D_ estimates for OT-I TCR binding higher-affinity pMHC, inactive pMHC would appreciably affect K_D_ estimates for lower-affinity pMHCs (Fig. 2H). As expected, a larger distortion would be observed with a higher fraction of inactive pMHC (Fig. S4).

Using the apparent K_D_ and the fraction of active pMHC, we were able to estimated the K_D_ of OT-I TCR binding to active pMHC (Fig. 2I). To validate this new method, we used it to measure the affinity of OT-I binding to VSV pMHC and two different levels of partially-denatured VSV pMHC (Fig. S5). The apparent K_D_, estimated using the original method, displayed wide variation across these three surfaces, while on the other hand, our new method produced the same active K_D_ value on all surfaces despite variations in the amount of denatured pMHC. This confirmed that our method can reliably estimate active K_D_ values. When we applied this extended workflow to estimate the active K_D_ for all pMHC, we found that it produced similar values for higher-affinity interactions but up to 4-fold higher K_D_ values for the lower-affinity interactions (Fig. 2J, Table 1).

### Revised OT-I affinities display much larger variation and weaker binding to self pMHC

The revised affinities that we now report for OT-I TCR binding various pMHCs at 37*^◦^*C are very different from the values previously reported, and the discrepancies are most pronounced for low-affinity peptides (Fig. 3A). For example, the OT-I TCR was originally reported to bind E1 pMHC with a K_D_ of 26.8 *µ*M, whereas we now report a 160-fold lower affinity of 3637 *µ*M. Our measurements are in closer agreement with five affinities more recently measured at 25*^◦^*C (11) (Fig. 3B). Importantly, our revised OT-I/pMHC affinities show much greater variation than previously reported. For example, we find a 150-fold variation in K_D_ from 34 *µ*M to 5066 *µ*M for the N4 and R4 pMHC, respectively. In contrast, Alam *et al.* (3) described only a 2-fold difference between OT-I TCR affinities for N4 and R4 pMHCs.

**Figure 3:**
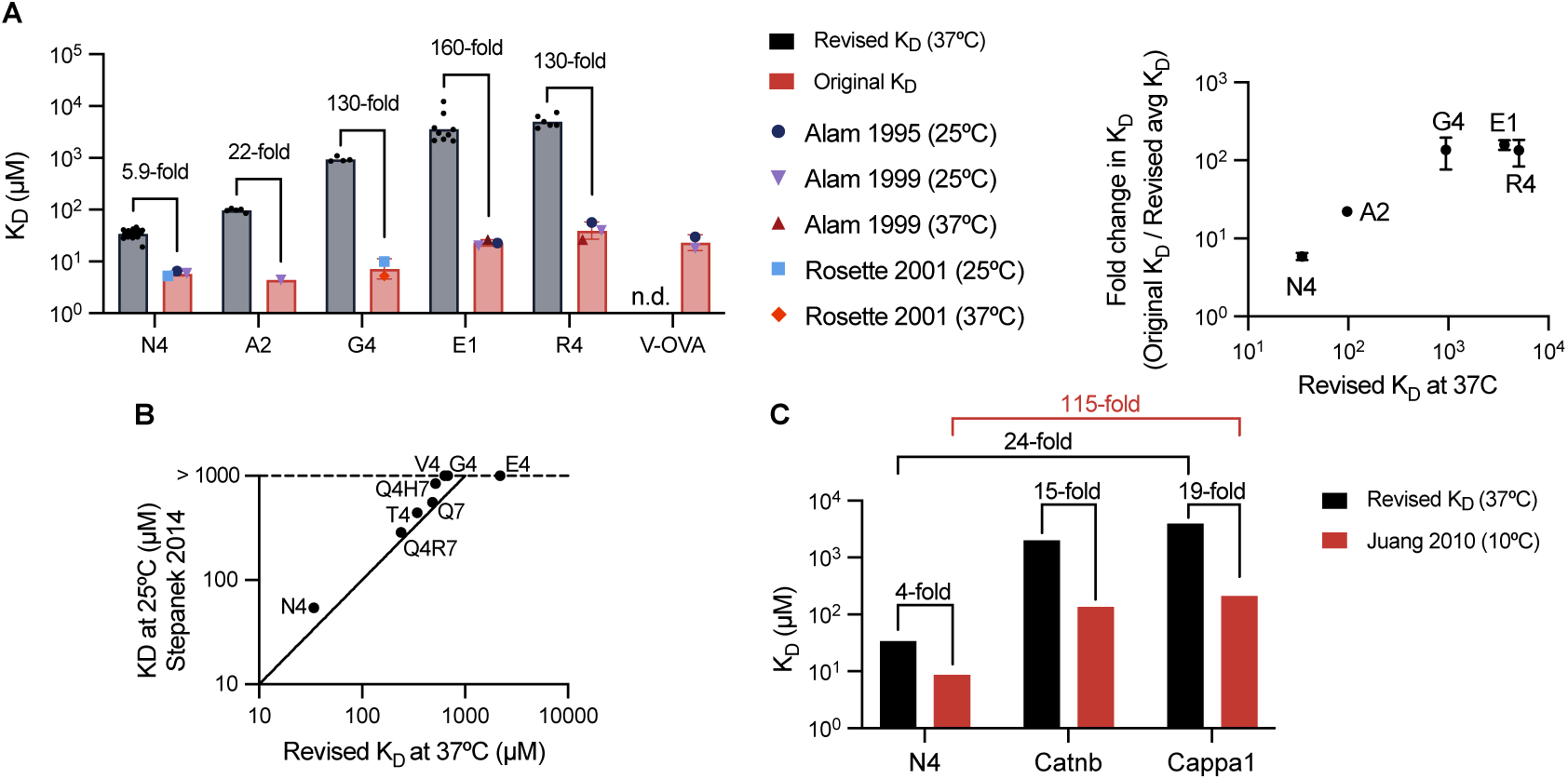
Discrepancies between published and revised OT-I affinities. **(A)** Comparison between original OT-I K_D_ values measured by SPR at 25*^◦^*C and 37*^◦^*C (2, 3, 5) and the revised K_D_ values measured at 37*^◦^*C in the present work as bar graphs (left) or scatter plot of fold-change (right). The original measurements at 37*^◦^*C for N4 and A2 were excluded because they displayed biphasic binding making affinity and kinetic estimates unreliable. Bar graphs show mean K_D_ values. **(B)** Comparison of K_D_ values determined at 25*^◦^*C (11) with our revised K_D_ values at 37*^◦^*C. Solid line indicates the identity line (y=x). **(C)** Comparison of K_D_ values measured for self-peptides at 10 *^◦^*C (10) and our revised K_D_ values at 37*^◦^*C.

Although self peptides have been identified for the OT-I TCR, previous estimates of their affinities have only been performed at the unphysiologically low temperature of 10*^◦^*C (10). Using our method, we show that the OT-I TCR binds Catnb and Cappa1 self peptides with K_D_ values of 2024 *µ*M and 3956 *µ*M, respectively, at 37 *^◦^*C, which are appreciably larger than the K_D_ values of 136 *µ*M and 211 *µ*M reported at 10*^◦^*C (Fig. 3C). Whereas the K_D_ varied by 24-fold between the foreign (N4) and self (Cappa1) antigens at 10*^◦^*C, we now find a much larger variation of 115-fold at 37 *^◦^*C. This suggests that thymic positive selection can proceed with ultra-low affinities enabling a much larger affinity window between foreign and self antigens in the periphery.

### Revised 3D affinities correlate well with 2D affinities

The process of antigen recognition takes place at the T cell - APC contact interface where both TCRs and pMHCs are attached to membranes that confine their movements to two-dimensions. It has long been speculated that 2D TCR/pMHC binding properties may not correlate well with 3D properties because of the impact of molecular forces on receptor/ligand interactions at membrane interfaces (37). In apparent support of this, original OT-I TCR/pMHC 3D affinity values did not correlated well with 2D affinity values, displaying a highly non-linear power relationship with a power (or slope on log-transformed values) of 2.9 (Fig. 4A). In other words, small changes in 3D affinity were associated with large changes in 2D affinity. This suggested that the impact of force on TCR/pMHC interactions may improve antigen discrimination (20). In contrast, our revised 3D affinity values correlate well with 2D affinity values with a much shallower slope of 1.3 (Fig. 4B,C). These new findings are similar to those reported for the 1E6 TCR, where the 3D and 2D affinities correlate very well with a slope of 1.0 (Fig. S7) (39). The finding that 3D affinity produces a high linear correlation with the 2D affinity for the OT-I and 1E6 TCRs argues against a substantial role for force in 2D TCR/pMHC binding in these settings.

**Figure 4:**
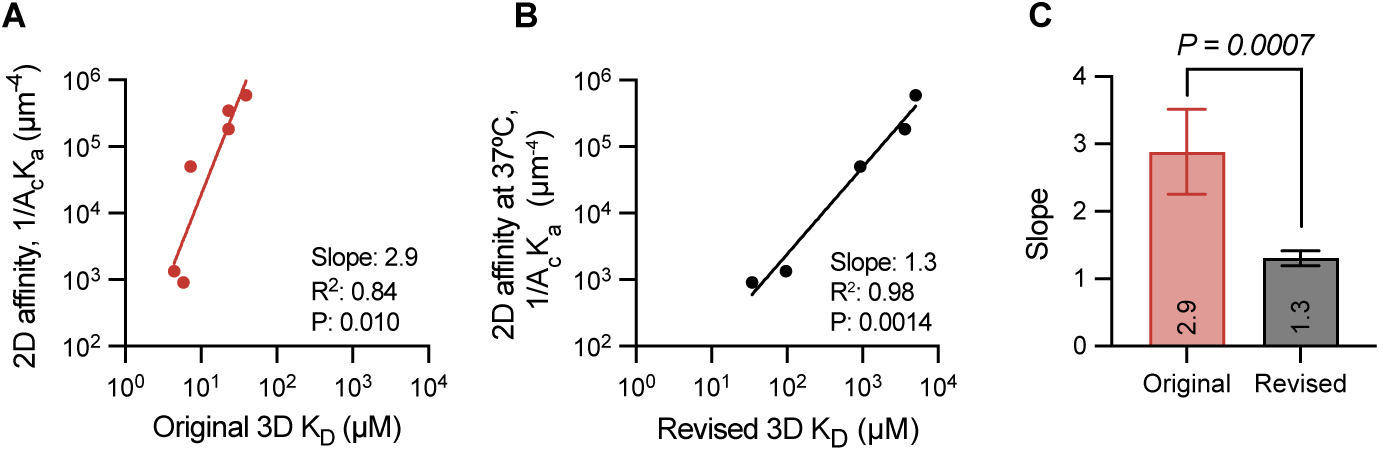
Revised 3D affinities produce high correlations with 2D affinities and display similar variation. (A-B) Correlation between 2D affinity values (20) with the (A) original and (B) our revised 3D K_D_ values. **(C)** Fitted slopes and standard errors from the fitted lines in A,B. An F-test determines the p-value for the null hypothesis that a single slope can fit both correlations.

### The OT-I TCR displays enhanced but imperfect antigen discrimination

Given the large differences between the originally reported OT-I affinities and the revised ones reported here, we measured the potency of these peptides in functional assays. Using naive CD8^+^ T cells from OT-I TCR transgenic mice, we quantified T cell activation potency of 8 different peptides (Fig. 5A). There was a significant correlation between peptide potency (EC_50_) and our revised K_D_ values but not with the original K_D_ values (Fig. 5B). Another key difference was the discriminatory power which is obtained by the slope of the relationship. The original K_D_ measurements produced a steep slope (*α* = 16). This indicates that small reduction in OT-I/pMHC affinity would abolish the T cell response, which has been termed near-perfect or absolute discrimination (19, 21, 34). In contrast, our revised K_D_ measurements produced a more modest discriminatory power (*α* = 2.4), representing what we have previously termed enhanced but imperfect antigen discrimination (38).

**Figure 5:**
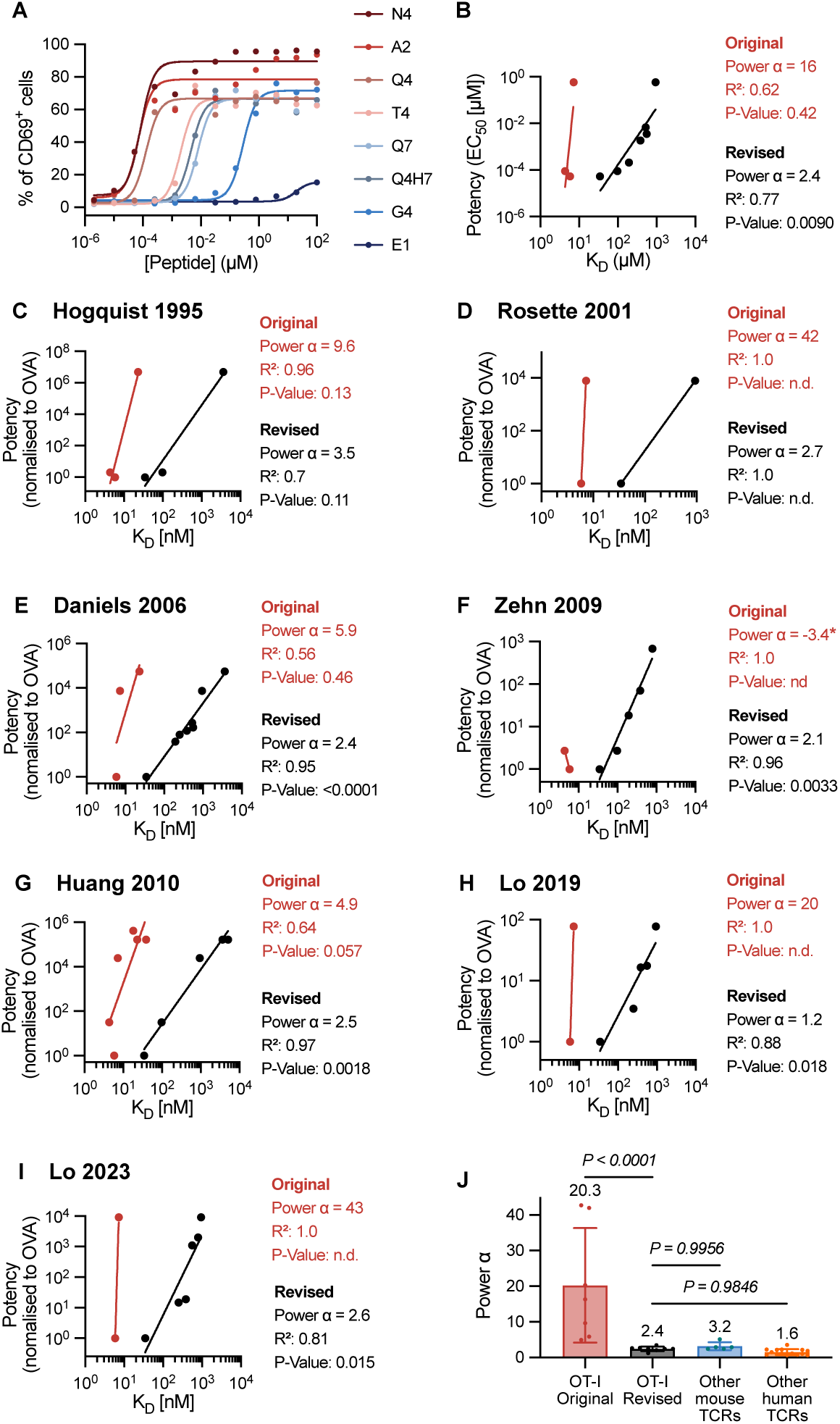
Revised 3D affinities correlate with OT-I T cell responses and reveal enhanced but imperfect antigen discrimination. (A,B) Representative OT-I T cell activation by the indicated peptides (A) and peptide potency (EC_50_) over K_D_ (B) for N=2 independent experiments. **(C-I)** Published potency data from the indicated study over original or revised K_D_ values. A power law (potency ∼ (K_D_)*^α^*) is fit to the data to estimate the discriminatory power (*α*). A Pearson correlation is used to determine *R*^2^ and p-values on log-transformed values. **(J)** The discriminatory power from panels B-I (OT-I original and revised) and from other mouse and human TCRs (see Pettmann et al (38)). Negative values for *α* as observed for (F) were excluded. A one-way ANOVA determines p-values. The original K_D_ values are the average K_D_ values from Fig. 3A.

We next plotted the potency data from 7 previous functional studies against our revised K_D_ measurements (1, 5, 8, 20, 25, 46, 47) (Fig. 5C-I). We found that, whereas the original K_D_ measurements correlated poorly with potency, and indicated a near-perfect discriminatory power of ∼20.3, our revised K_D_ measurements correlated well with potency, and indicated enhanced but imperfect discriminatory power of ∼2.4 (Fig. 5J). This conclusion holds when discriminatory power is calculated using apparent K_D_ values, confirming that our method to determine active K_D_ values does not affect our conclusion (Fig. S6).

The self-peptides Catnb and Cappa1 have recently been shown to activate OT-I T cells at very high concentrations (∼100 *µ*M) (25). This is consistent with their very low affinities (Fig. 3J). Indeed, like Catnb and Cappa1, other very low affinity peptides such as E1 and R4 have also been shown to induce positive selection (1, 48). This suggests that, while T cells can maintain tolerance to self pMHCs (K_D_ *>*2000 *µ*M) when expressed at normal levels, this tolerance can be broken if these self peptides are abnormally over-expressed.

### The discriminatory power of the OT-I TCR increases with K_D_, consistent with the kinetic proofreading mechanism

The discriminatory power is an empirical measure of antigen discrimination and a mechanistic description is provided by the kinetic proofreading model (Fig. 6A). In this model, a time-delay (*τ*_kp_) between pMHC binding and TCR signalling produced by a series of biochemical signalling steps (N, each with rate *k*_p_) reduces the probability of productive signaling by pMHCs with a short dwell time. To estimate these parameters, we fit the model to the averaged potency of each OT-I ligand using our revised 3D affinity measurements (Fig. 6B). Although the number of proofreading steps for the OT-I TCR (N=4.2) is larger than previous reports for the 1G4 TCR and a CAR (N*<*3) (38, 49), the overall time-delay is shorter (*τ*_kp_ = 0.21 s for OT-I vs 2.7 s for 1G4) because of a much higher rate of traversing each step (21 s*^−^*^1^ for OT-I vs 1.0 s*^−^*^1^ for the 1G4). These fitted proofreading parameters that explain a discriminatory power of 2.4 are also consistent with the high antigen sensitivity of the TCR (Fig. 6C).

**Figure 6:**
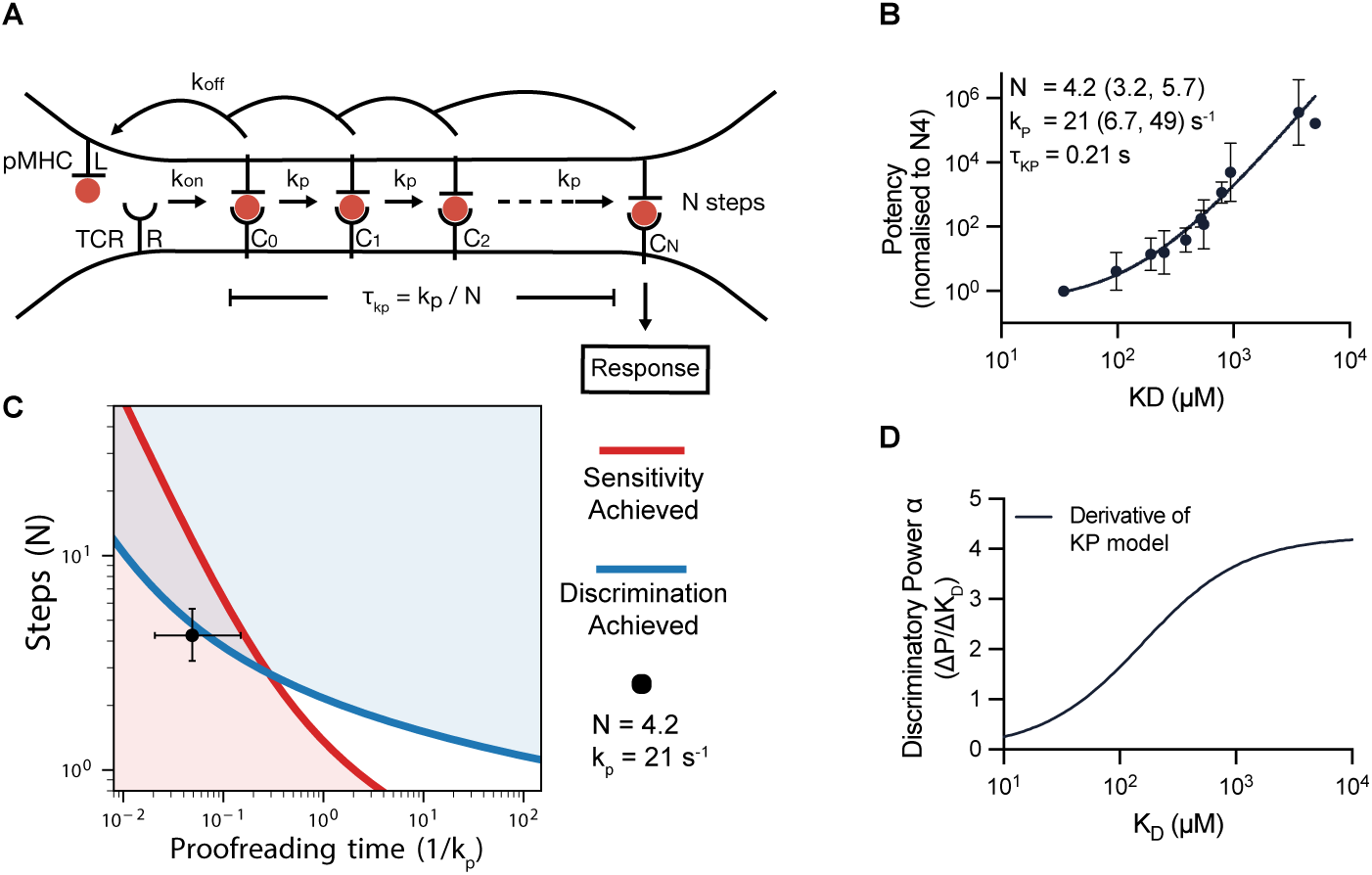
The kinetic proofreading model explains antigen discrimination by OT-I T cells with a short proofreading time-delay and highlights that the discriminatory power can change with affinity. **(A)** Schematic of the kinetic proofreading model. **(B)** Potency over revised K_D_ values fitted by the kinetic proofreading model (solid line). Potency data from all studies shown in Fig. 5 is normalised to N4 within each study before averaging across all studies and displayed as mean±SD. The mean and 95%CI of the best-fit parameters are shown. **(C)** A binary heatmap displaying regions that achieve a discriminatory power of 2.4 (blue) and high antigen sensitivity (red). The fitted number of steps (N) and proofreading time (1/*k_p_*) along with 95% CI is shown as a dot with error bars, which overlap with a region where both discrimination and sensitivity are achieved. The binary heatmap is produced as described in Petttmann et al (38) with the off-rate estimated as *k*_off_ = K_D_ · *k*_on_, where K_D_ =34 *µ*M and *k*_on_ = 0.13 µM*^−^*^1^s*^−^*^1^ (for N4) and all other parameters as in Pettmann *et al.* (38). **(D)** The discriminatory power (rate of change of potency with respect to K_D_) over K_D_ determined by taking the first derivative of the solid line in (B).

We noted that when plotting all the potency data against affinity a non-linear relationship emerges (Fig. 6B). This indicates that the discriminatory power-law relationship between potency and affinity has a discriminatory power that varies with affinity. Indeed, a plot of discriminatory power against affinity shows that it increases from near 0 at low K_D_ to 4 at high K_D_ values (Fig. 6D). In other words, the discriminatory power of the OT-I TCR is highest for lower affinity antigens, a property predicted by the kinetic proofreading model.

## Discussion

Despite the fact that OT-I TCR transgenic mice are widely used, there was no accurate affinity data for the OT-I TCR binding relevant pMHCs at physiological temperatures, and original measurements showed unusual bi-phasic binding at 37*^◦^*C. These original measurements gave rise to the notion that the OT-I TCR displays near-perfect antigen discrimination based on affinity and that the 3D affinity measurements do not correlate with 2D affinity measurements. Here, we developed a new method to accurately measure ultra-low affinities and used it to systematically measure the OT-I TCR affinity to 19 commonly used peptides. We found that our revised 3D K_D_ values correlate well with 2D K_D_ values, and with T cell functional responses. Importantly, these K_D_ values, together with functional data from many laboratories, demonstrate that the OT-I TCR displays enhanced but imperfect antigen discrimination, as has been reported for other TCRs (38). Finally, we have shown that the discriminatory power of the OT-I TCR is highest for low affinity pMHC ligands, a result explained by the kinetic proof-reading model.

In control experiments we found that the OT-I TCR binds non-specifically to unfolded MHC and we estimated this affinity to be K_D_ ∼ 2000 *µ*M (Fig. 2). This binding may represent binding to the empty, relatively non-specific peptide-binding groove in unfolded pMHC. Consistent with this, it has been observed that a TCR can weakly bind empty MHC molecules but not those loaded with irrelevant peptides (50). When the fraction of inactive pMHC is small, their presence is unlikely to impact the accuracy of higher-affinity measurements. However, even a small fraction of inactive pMHC can be problematic when measuring very low affinities. We developed a simple workflow to quantify the amount of inactive pMHC enabling us to extract the K_D_ for active pMHC from the apparent K_D_ values. This conversion relied on an accurate estimate of the amount of inactive pMHC, which we estimated by injection of conformationally sensitive antibodies to pMHC at the end of each experiment. We demonstrated that we can extract the same active K_D_ values from OT-I binding to surfaces with different levels of inactive pMHC (Fig. S5). Interestingly, a loss of peptide and *β*_2_m from cell surface MHC class I has been suggested to induce homodimerization via the *α*3 domains, followed by internalization (51). This would reduce the likelihood that unfolded MHC on the APC would bind to TCRs or other receptors, activating cells non-specifically.

We found a linear correlation between the TCR/pMHC affinities measured using purified proteins in solution (i.e. 3D K_D_) and previously reported 2D affinities (Fig. 4). This contrasts with the highly nonlinear correlation between the 2D and the original reported 3D K_D_ values (20). A linear correlation between 3D and 2D affinities has also been reported for the 1E6 TCR (39) (Fig. S7). These results suggest that, despite differences between the TCR/pMHC interaction in solution and within cell-cell interfaces, T cells accurately measure linear proxies of the 3D K_D_. This is unexpected because T cells generate large mechanical forces during antigen recognition (52, 53) and molecular forces can have large non-linear effects on the TCR/pMHC bond lifetime (22, 54–57). These results can be reconciled by the force-shielding model (45), which proposes that T cells deploy mechanisms to shield the TCR/pMHC interaction from molecular forces. By eliminating force, the 2D TCR/pMHC lifetimes would be expected to correlate with the 3D lifetimes or 3D K_D_ (when *k*_on_ displays minimal variation). Consistent with this model, the 2D and 3D life-times have been shown to be similar (40) and most TCR/pMHC interactions appear to take place without experiencing forces (58). We have suggested that receptor/ligand interactions, such as CD2/CD58 and/or LFA-1/ICAM-1, mediate force-shielding and this would require spatial redistribution near the TCR/pMHC (45). This is supported by the observation that ligand mobility is required to abolish TCR/pMHC forces (59). Taken together, the high linear correlation between 3D and 2D K_D_ values suggests that, although T cells generate large mechanical forces during the process of antigen recognition, they deploy mechanisms to shield TCR/pMHC interactions from these forces.

The original K_D_ values reported for OT-I suggested that T cells possessed near-perfect discriminatory powers of ∼5-40 (Fig. 5J). For example, while the OT-I TCR bound the E1 peptide with a 3-fold lower affinity than the N4 peptide, OT-I T cells required a 100,000-fold higher concentration of E1 than N4 peptides to be activated. In contrast, other TCRs exhibit a discriminatory power of approximately 2, where T cells require only 3^2^ = 9-fold higher concentrations to be activated by peptide antigens with a 3-fold lower affinity (38). Our revised affinity measurements show that the OT-I TCR binds the E1 peptide with a 106-fold lower affinity than the N4 peptide (Fig. 2J), rather than the previously reported 3-fold lower affinity. Thus, OT-I T cells failed to respond to E1, not because of their exceptionally high discriminatory power, but because the OT-I TCR binds E1 with exceptionally low affinity. Using these K_D_ values we calculate the discriminatory power of OT-I TCR to be approximately 2.4, similar to other TCRs. Our new results therefore resolve this major discrepancy between the OT-I TCR and other TCRs.

Like other murine TCRs (discriminatory power 3.2), the OT-I TCR appears to have a greater discriminatory power than human TCRs (2.4 vs 2.0) (38). It also has different kinetic proof-reading parameters with a time-delay that is more than 10-fold shorter than the human 1G4 TCR (2.7 s compared to 0.21 s for OT-I) (38). A key difference between studies on murine and human TCRs is that the murine T cells, like the OT-I, are often obtained from TCR transgenic mice where they have undergone development while expressing the TCR. In contrast, human TCRs are often studied after transfection into polyclonal primary T cells. Since T cells undergo tuning during development based on the TCRs they express (60), this is likely to affect their sensitivity and discriminatory power. Although incompletely understood, one mechanism for tuning is altering expression surface CD5 levels (61), which we have shown modifies TCR discrimination (62). The fact that the OT-I has the shortest 3D half-life ever reported for a TCR (45), and the resulting tuning of signalling by OT-I TCR expressing T cells to adapt to this short half-life, may contribute to the difference between its kinetic proofreading parameters and those observed for the 1G4 TCR expressed in polyclonal human T cells.

The enhanced but imperfect discriminatory power of the OT-I TCR and the ultra-low affinities that we now report for self antigens have two important implications for peripheral tolerance. First, it suggests that ultra-low affinity self-antigens can induce thymic positive selection and as a result, the affinity window between foreign and self antigens is likely much larger than previously believed (Fig. 3C). Second, it suggests that peripheral tolerance can be broken by self pMHCs provided they are expressed at sufficiently high concentrations. Indeed, the self peptides Catnb and Cappa1 can activate OT-I T cells when presented at ∼ 10^5^-fold higher concentrations compared to the foreign antigen N4 (10 vs 10*^−^*^4^ *µ*M) (25). This is consistent with an imperfect discriminatory power of 2.4 that would predict T cell responses by these self peptides when their concentration increases by ∼ 100^2.4^ = 63, 000. This supports the notion that, in addition to recognising foreign antigens, T cells have a homeostatic surveillance role in detecting cells expressing aberrantly high levels of protein, such as hormone-secreting endocrine tumours (63).

One prediction of the kinetic-proofreading model is that the discriminatory power of a TCR should increase with K_D_, i.e. discrimination is largest for lower-affinity pMHCs (38). This prediction has been difficult to test in the absence of sufficient TCR/pMHCs measurements over a sufficiently wide range of K_D_ values. The large number of OT-I functional studies using pMHCs whose affinities we now report to have a wide-range of K_D_ values has enabled us to accurately measure how the discriminatory power varies with K_D_ for the first time (Fig. 6D). This has highlighted that the discriminatory power can be as high as ∼4 at the ultra-low affinity range whereas it decrease to values below 1 at the higher-affinity range where potency begins to saturate. Additional studies are required to establish whether this is a general feature of TCR discrimination.

In conclusion, by accurately measuring 3D K_D_ values between the OT-I and 19 pMHCs, including foreign and self antigens, we have reconciled reported discrepancies between the OT-I TCR and other TCRs, confirmed that the OT-I displays physiological affinity to its foreign antigen, shown that the 3D affinity predicts the 2D affinity, and that the OT-I TCR displays enhanced but imperfect discrimination, which increases for lower-affinity antigens. Collectively, these results highlight that despite the complex and mechanically active T cell/APC interface, T cells make decisions based on proxies for the 3D K_D_ measured with purified proteins in solution.

## Materials & Methods

### Protein expression and purification

#### OT-I TCR

For affinity measurements with soluble TCR, we used an OT-I TCR construct consisting of the murine variable OT-I domain and the human constant domain truncated above the transmembrane domain with an artificial interchain disulphide, as described previously (11). TCR *α* and *β* chains were expressed in BL21 DE3 *Escherichia coli* cells following induction with 0.15 mM IPTG and isolated from inclusion bodies. Proteins were stored at-80*^◦^*C until use.

OT-I-TCR was refolded by adding 15 mg of each chain dropwise in 1 L refolding buffer (150 mM Tris-HCl (pH 8.0), 3 M Urea, 200 mM Arg-HCl, 0.5 mM EDTA, 0.1 mM PMSF), followed by dialysis for 3 days in 10 L Tris buffer (10 mM Tris-HCl (pH 8.5)), with a buffer change after 24 h. After dialysis, the protein was filtered and purified using ion-exchange chromatography (HiTrap Q column [Cytiva]) with a NaCl gradient in the dialysis buffer. Next, protein was concentrated and purified again by size exclusion chromatography (Superdex 200 Increase column [Cytiva]) in HBS-EP buffer (0.01 M HEPES pH 7.4, 0.15 M NaCl, 3 mM EDTA, 0.005% v/v Tween-20). Purified TCR was used for SPR measurements not longer than 24 hours after purification to avoid aggregation. Protein concentration was measured with Nanodrop.

#### pMHCs

Class I pMHCs were generated using mouse H-2K^b^ heavy chain and human beta-2 microglobulin (*β*2m), biotinylated on the C terminus of the heavy chain. pMHCs produced in HEK293T cells were biotinylated and peptide-exchanged by the NIH tetramer facility. For pMHC produced in *E.coli*, soluble mouse H-2K^b^ heavy chain with a C-terminal AviTag/BirA recognition sequence and human *β*-2m were expressed separately in BL21 DE3 *E. coli* cells and isolated from inclusion bodies. MHC heavy chain, *β*-2m and peptide were then added dropwise to the refolding buffer (100 mM Tris-HCl, pH 8.0, 400 mM L-Arg·HCl, 2 mM EDTA, 5 mM Reduced glutathione, 0.5 mM Oxidised glutathione, 0.1 mM PMSF) at a concentration of 2 *µ*M, 1 *µ*M, 10 *µ*M, respectively. The protein solution was kept under constant stirring for 48 h at 4*^◦^*C. Afterwards, the refold was filtered through a 0.45 µL filter and concentrated using centrifugal filters. pMHCs were biotinylated overnight at room temperature using the BirA Biotin-protein ligase bulk reaction kit (Avidity LLC). Next, pMHCs were purified by size exclusion chromatography (Superdex 75 column [GE Healthcare]) in HBS-EP Buffer. pMHCs were aliquoted and stored at-80*^◦^*C until use.

#### Surface plasmon resonance

Affinities of the OT-I TCR to peptide variants were measured with a newly established SPR technique (SPR) for ultra-low TCR-pMHC affinities described previously (38). We generated soluble OT-I TCR, which recognises the ovalbumin (OVA) peptide (SIINFEKL) loaded onto a murine H-2K^b^ class I MHC. Equilibrium binding analysis of TCR-pMHC interactions was performed by SPR on a Biacore T200 instrument (GE Healthcare Life Sciences) with CM5 sensor chips. HBS-EP was used as running buffer and all K_D_ measurements were performed at 37°C. For protein immobilisation, the sensor chip was saturated with streptavidin using an amino coupling kit (Cytiva). Biotinylated pMHCs were injected into experimental flow cells (FC) for different durations to immobilise 400 to 1500 RU pMHC. Matching levels of CD86 were immobilised in FC1 as a reference. Next, excess streptavidin was blocked with two 40 s injections of 500*µ*M biotin (Avidity) and the sensor was conditioned with 8 injections of running buffer. TCR was injected at increasing concentrations at 30 µl/min. Buffer was injected after every 2 or 3 TCR injections. Following TCR injections, anti-*β*2m antibody (B2M-01 Invitrogen, MA1-19141) that binds correctly folded pMHC was injected for 8 min at 10 µl/min.

#### Obtaining K_D_ (apparent) from SPR data

Apparent K_D_ values were obtained by fitting a 1:1 binding model (RUeq = B_max_ ·[TCR]/(K_D_ +[TCR])) to the double referenced equilibrium RU values. For low affinity antigens, this curve does not saturate at the highest TCR concentration, therefore an accurate prediction of B_max_ and thus K_D_ is not possible. Instead, the high affinity the N4 pMHC-TCR interaction was used to generate the empirical standard curve to relate the B_max_ of TCR binding to the maximal antibody binding. For low affinity peptides, B_max_ was constrained to B_max_ inferred from the standard curve when fitting the SPR data.

#### Simulation of TCR Binding to mixed populations of active and inactive pMHC

Steady-state binding response of the TCR interacting with a mixture of active (correctly-folded) and inactive pMHC was modeled using the following equation:

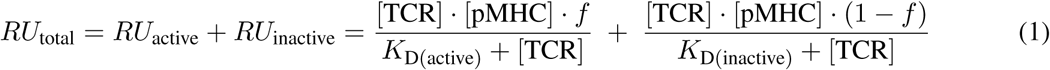

Here, *f* is the fraction of active pMHC, [TCR] and [pMHC] are the concentrations of TCR and pMHC, respectively, *K*_D(active)_ is the dissociation constant for the active pMHC-TCR interaction, and *K*_D(inactive)_ is the dissociation constant for the inactive pMHC-TCR interaction.

#### Determination of the fraction of active pMHC

The fraction of active pMHC (*f*) was determined from SPR measurements by comparing the TCR B_max_ to the pMHC immobilisation level. The TCR B_max_ was obtained using the B2M-01 antibody and the empirical standard curve in Fig. 1B (as described above). Thus,

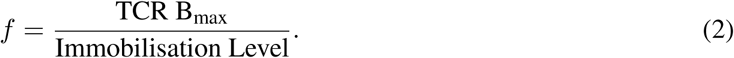

#### Fitting SPR experimental data to obtain K_D(active)_

SPR experimental data were fitted with Eq. 1, with *K*_D(inactive)_ = 2180 *µ*M and *f* determined using Eq. 2. This procedure provided the K_D_ value for the active fraction (*K*_D(active)_).

#### Interpolating K_D(active)_ from K_D(apparent)_

To relate *K*_D(active)_ to *K*_D(apparent)_, simulated binding curves were generated for a range of *K*_D(active)_ values at a given active fraction *f* using Eq. 1. These simulated datasets were then fitted with a 1:1 bind-ing model (RUeq = B_max_ ·[TCR]/(K_D_ +[TCR])) to obtain *K*_D(apparent)_. The resulting relationship between *K*_D(active)_ and *K*_D(apparent)_ was used to interpolate *K*_D(active)_ for all peptides, given their experimentally determined *K*_D(apparent)_ (Fig. S1 and their average *f* (Fig. 2F).

#### Exclusion Criteria

If a given *K*_D(apparent)_ fell outside the range of the *K*_D(active)_-to-*K*_D(apparent)_ curve (i.e., beyond the curve’s saturation point), the data point was excluded. Under these circumstances, the observed TCR binding could be entirely explained by interactions with inactive pMHC alone. Applying this exclusion criterion removed one data point for peptide E1 from the final analysis.

## Mice

OT-I mice (JAX stock no.: 003831) were purchased from Jackson Laboratory and CD45.1 mice from Charles River. Mice were bred and maintained in the University of Oxford specific pathogen-free (SPF) animal facilities. Mice were routinely screened for the absence of pathogens and were kept in individually ventilated cages with environmental enrichment at 20–24*^◦^*C, 45–65% humidity with a 12 h light/dark cycle (7am–7pm) with half an hour dawn and dusk period. Mice were euthanized by CO2 asphyxiation followed by cervical dissociation. Breeding was conducted in agreement with the United Kingdom Animal Scientific Procedures Act of 1986 and performed under approved experimental procedures by the Home Office and the Local Ethics Reviews Committee (University of Oxford) under UK project licenses P4BEAEBB5 and PP3609558.

## T cell activation

OT-I T-cells were isolated from lymph nodes and spleen of 6-12 week old OT-I mice. Selection was carried out with a MojoSortTM CD8+ T-cell negative isolation kit and magnets (Biolegend, #480008 and #480019). Isolated OT-I T-cells were resuspended in complete RPMI (RPMI 1640 [Gibco, #21870-076] supplemented with 2% FCS and 100x Penicillin-Streptomycin [Gibco, #10378-016]). Naive OT-I T-cells (50,000) were seeded in 96-well U-bottom together with splenocytes (100,000) from CD45.1 mice loaded with the indicated dose of the following peptides: N4 (SIINFEKL), A2 (SAINFEKL), Q4 (SIIQFEKL), T4 (SIITFEKL), Q7 (SIINFEQL), Q4H7 (SIIQFEHL), G4 (SIIGFEKL), E1 (EIINFEKL). Cells were harvested after 24 hours.

## Flow Cytometry

Single-cell suspensions obtained from spleen or cultured CD8+ T-cells were stained in V-bottom 96-well plates in flow cytometry buffer (2% FCS, 2 mM EDTA, and 0.02% sodium azide in 1x PBS). Live dead staining and surface staining was performed using Zombie NIR Fixable Viability Kit (Biolegend, #423106/423105), TruStain FcXTM (anti-mouse CD16/32, Biolegend, #101319) and fluorochrome-conjugated the primary antibodies against CD45.1 (Biolegend, clone: A20), CD8 (Biolegend, clone: 53-6.7), CD69 (Biolegend, clone: H1.2F3), and CD44 (Biolegend, clone: IM7). Cells were fixed using 4% PFA for 30 minutes at 4*^◦^*C. Flow cytometry data were recorded on BD LSRII or FortessaX20 using BDFACSDiva (v8.0) software and analyzed using FlowJoTM software (v10.4.2, Tree Star).

## Data Analysis

All data fitting and statistical analysis was carried out in GraphPad Prism 10.

### Obtaining antigen potency

The dose-response data from the T cell activation experiments was fitted with a four-parameter sigmoidal model in GraphPad Prism 10, using Least-Squares regression. The model was defined as:

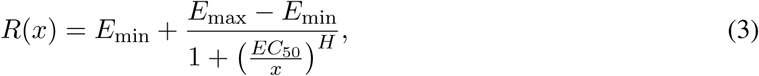

where *x* represents the peptide concentration used to load the antigen presenting cells (in µM). The fitted EC_50_ values were used as potency (*P*) values. EC_50_ values exceeding the highest tested peotide concentration were excluded, ensuring that no extrapolated results were used in the final analysis.

### Determination of discrimination power α

We obtained the discrimination power *α* by fitting a power law in log-space to the log-transformed potency over affinity data:

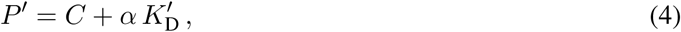

where *P ^′^* = log_10_(*P*) and *K^′^*= log_10_(*K_D_*).

### Fitting of the kinetic proofreading model

The log-transformed potency over affinity data was fit to the kinetic proofreading model using the following equation (38):

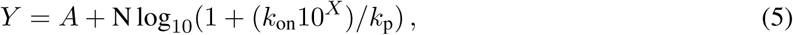

where *Y* is the log-transformed potency, *X* is the log-transformed affinity, *k*_on_ is the on-rate, *k*_p_ is the proofreading rate, N is the number of steps, and *A* is the maximum potency (y-intercept). Given that on-rates produce only modest variation between pMHCs, we fixed *k*_on_ to the value measured for the OT-I/N4 interaction (0.13 *µ*M*^−^*^1^s*^−^*^1^) (45).

## Data Availability

All data supporting the findings of this study are available within the paper and its Supplementary Information

## Acknowledgements

We thank Andrew Sewell and David Cole for providing plasmids for OT-I TCR expression and the NIH tetramer facility for providing purified pMHC.

## Open access

This research was funded in whole, or in part, by the Wellcome Trust [207537/Z/17/Z]. For the purpose of Open Access, the author has applied a CC BY public copyright licence to any Author Accepted Manuscript version arising from this submission.

## Author contributions

Conceptualization (OD, AH, PAvdM), Data Curation (AH, MK), Formal Analysis (AH), Funding Acquisition (OD), Investigation (AH, MK, KM, LFJU, JMM, AG, OD), Methodology (AH, MK, AG, PAvDM, OD), Project Administration (OD), Supervision (AG, PAvdM, OD), Visualization (AH, OD), Writing – Original Draft (AH, OD, PAvdM), Writing – Review & Editing (AH, AG, MK, PAvdM, OD)

## Supplementary Information

**Figure S1:**
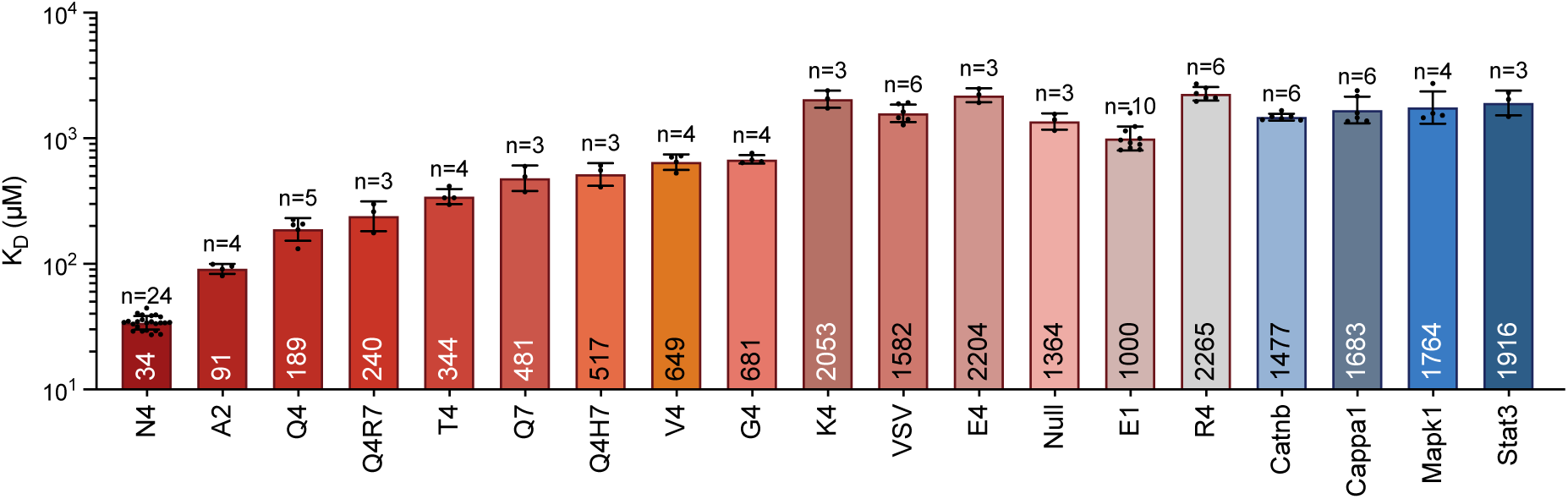
Apparent K_D_ values for OT-I specific peptides. The K_D_ values were obtained by fitting the steady-state binding using a 1:1 model with B_max_ constrained to value obtained from the B2M or Y3 antibody binding using standard curve (see Fig. 1 and Methods). Geometric mean is displayed within the bars, mean values, error and N are listed in Table 1.

**Figure S2:**
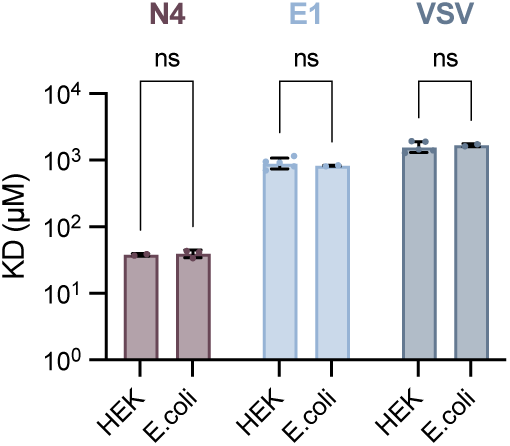
The OT-I TCR displays similar affinity to pMHC produced in *E.coli* and HEK293T cells. The *E.coli* produced pMHC where produced in-house whereas the HEK293T produced pMHC were supplied by the NIH tetramer facility.

**Figure S3:**
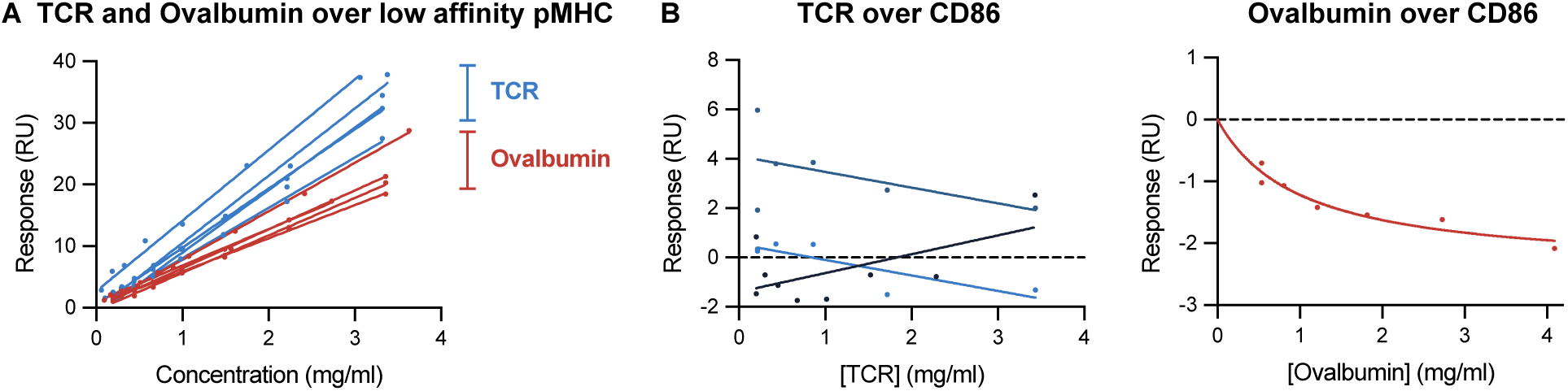
An irrelevant protein (ovalbumin) can bind heterotrimic pMHC but not monomeric CD86. **(A)** Steady-state binding of ovalbumin or OT-I analytes over pMHC surfaces. Note that the TCR displays higher binding than Ovalbumin suggesting that it binds both inactive and active pMHC whereas Ovalbumin binds only inactive pMHC. **(B)** Steady-state binding of TCR (left) or Ovalbumin (right) over CD86 surfaces reveals no detectable binding. Negative values can arise from modest refractive index effects.

**Figure S4:**
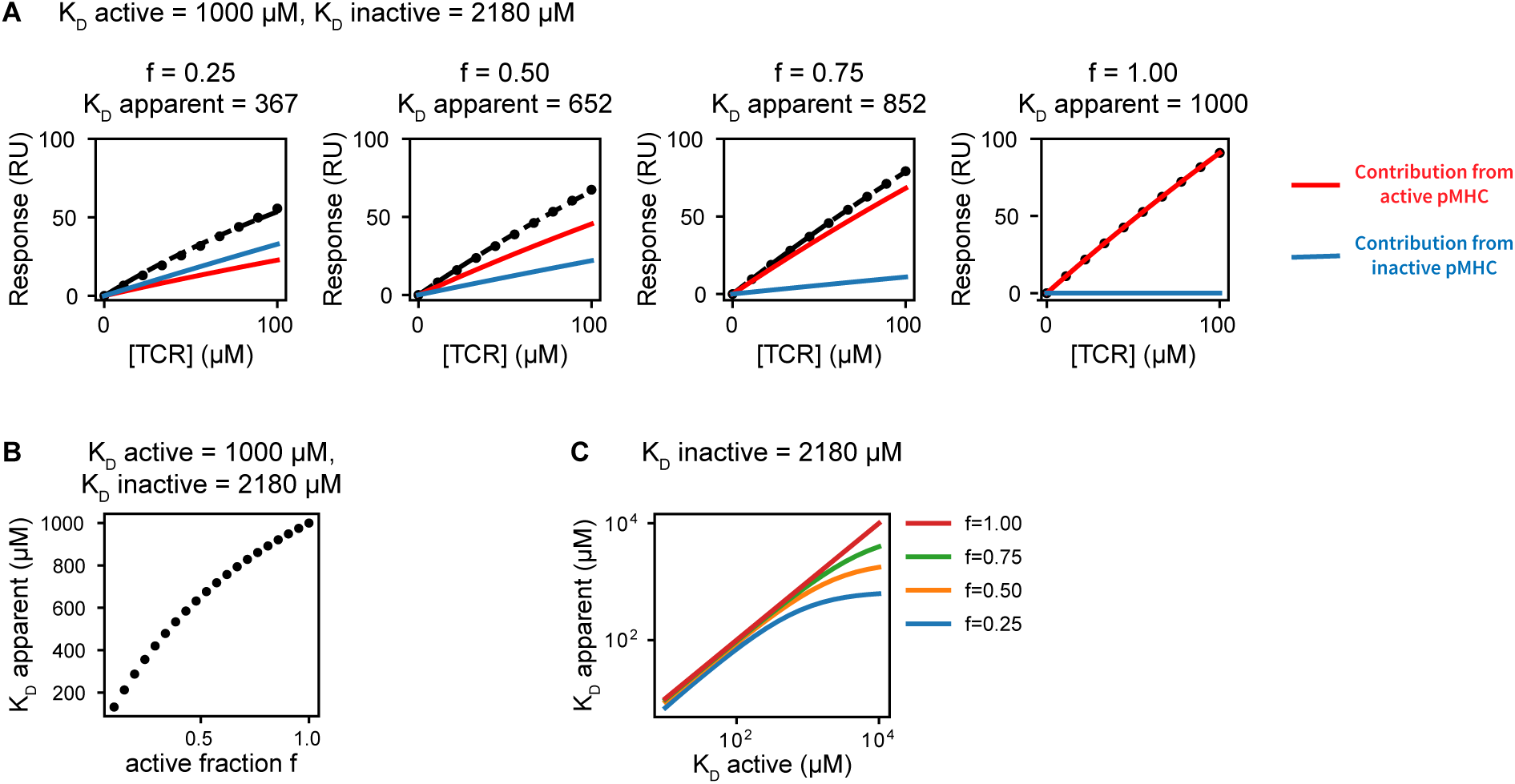
Simulations highlight the impact of the fraction of active pMHC (f) on apparent K_D_ values. **(A)** Simulated TCR binding curves to surfaces containing variable amount of active and inactive pMHCs (columns). The overall binding signal (black) was fit to a 1:1 binding model to estimate the apparent K_D_. **(B)** Apparent K_D_ values derived from fitting the simulated curves in (A) plotted against the fraction of active pMHC (f). The apparent K_D_ diverges from the active K_D_ as f decreases. **(C)** Apparent K_D_ values plotted against the true active K_D_ for different f, demonstrating the dependency of apparent K_D_ on both f and the active K_D_.

**Figure S5:**
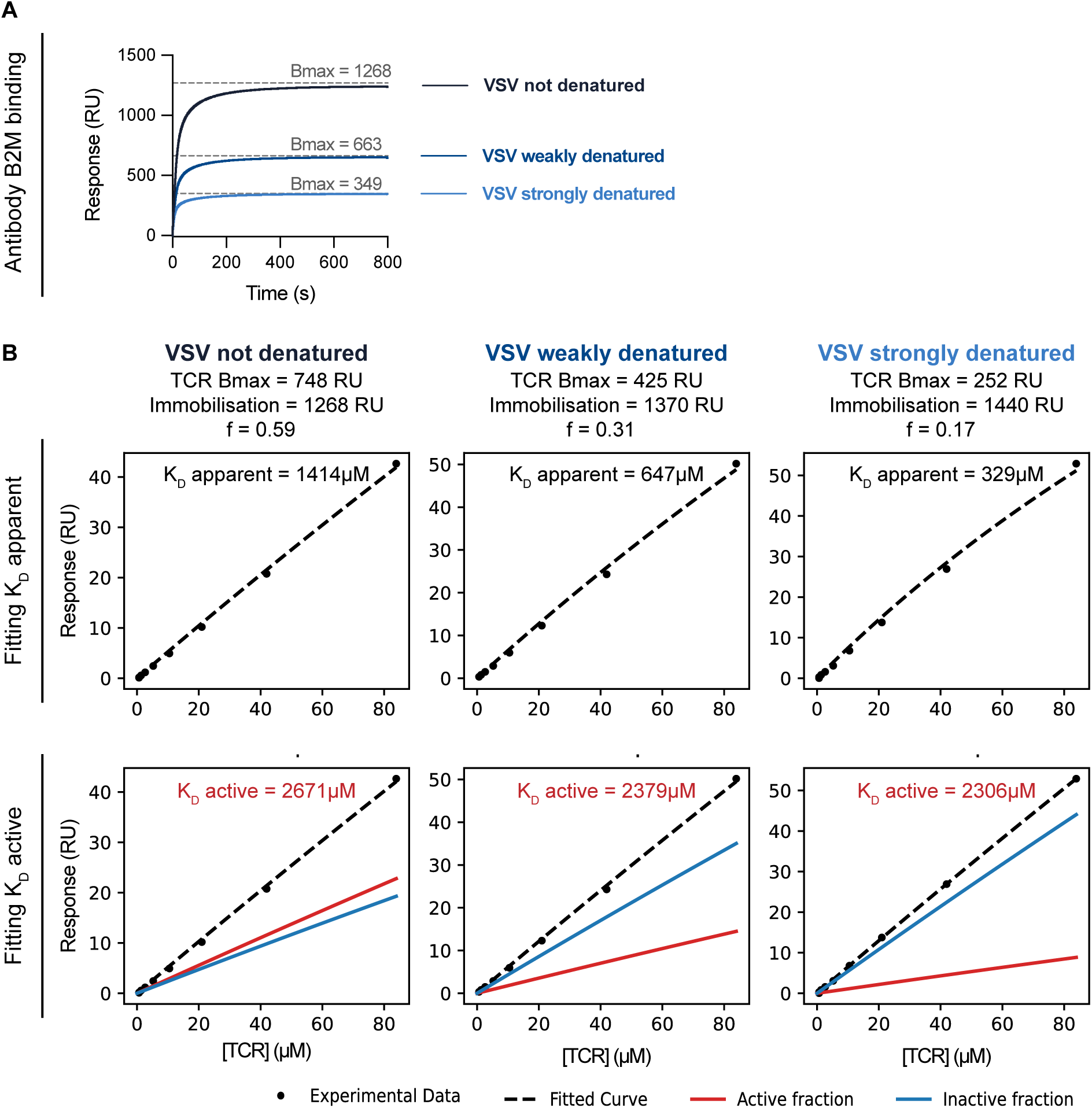
Calculating the active K_D_ provides similar results for OT-I binding to surfaces with large differences in the fraction of active pMHC. The VSV pMHC was immobilised at similar levels on 3 flow cells in SPR before inducing denaturing by a short (weakly denatured) or long (strongly denatured) injection of glycine solution (pH 1.7). **A** B2M antibody binding curves to the three VSV pMHC surfaces. The TCR B_max_ was estimated using the standard curve in Fig. 1B. **B** Steady-state TCR binding response to the 3 surfaces (columns). To determine the apparent K_D_, the data was fit with a 1:1 binding model with constrained B_max_ to determine K_D_ apparent (top row). To determine the active K_D_, the workflow in Fig. 2G-I was used, where the fraction of active pMHC was calculated from the ratio of TCR B_max_ to pMHC immobilisation, and K_D_ inactive was fixed to 2180 *µ*M. While the apparent K_D_ displayed large differences, the active K_D_ produced consistent results across all VSV pMHC surfaces.

**Figure S6:**
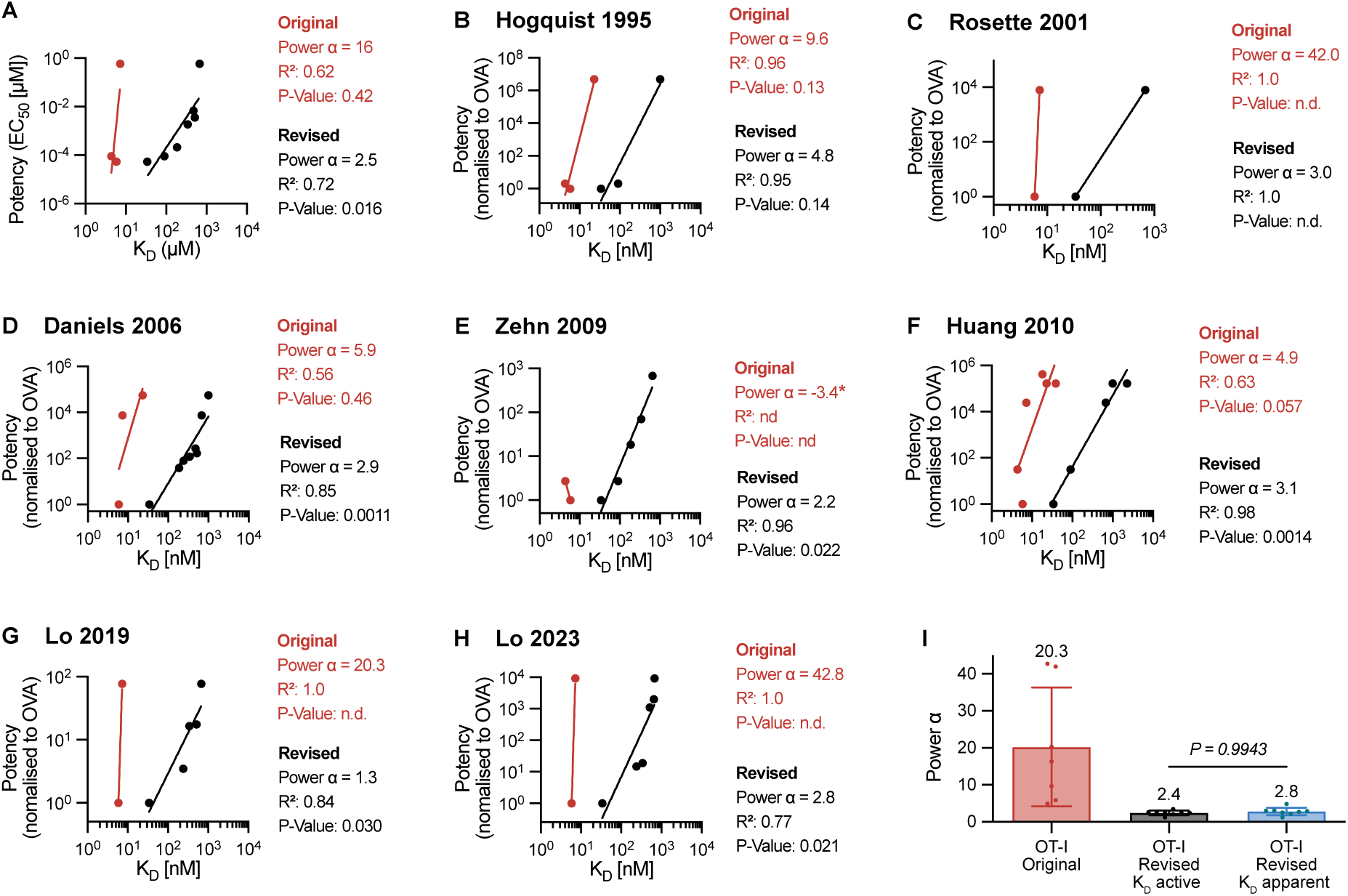
Discriminatory power of OT-I TCR calculated with K_D_ apparent shows imperfect discrimination. Plots show peptide potency over original or apparent K_D_ values. **(A)** Potency data from Fig 5A. Data is mean of N=2 independent experiments. **(B-H)** Published potency data from the indicated study over original or apparent K_D_ values. A power law (potency ∼ (K_D_)*^α^*) is fit to the data to estimate the discriminatory power *α*. A Pearson correlation is used to determine *R*^2^ and p-values on log-transformed values. **(I)** The discriminatory power from panels A-H in comparison with discriminatory power calculated with active K_D_ values. The p-value is determined using a t-test.

**Figure S7:**
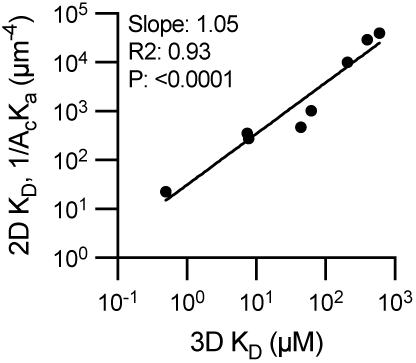
Quantitative comparison of 2D and 3D affinties for the 1E6 TCR. The log-transformed data was fitted with a linear regression. An F-test was used to determine a p-value for the null hypothesis that the slope is equal to zero. All data was taken from Cole et al (39).

