## Supplementary Information for "The 3D affinities of the OT-I TCR to foreign and self-antigens predict their 2D affinities and reveal imperfect antigen discrimination"

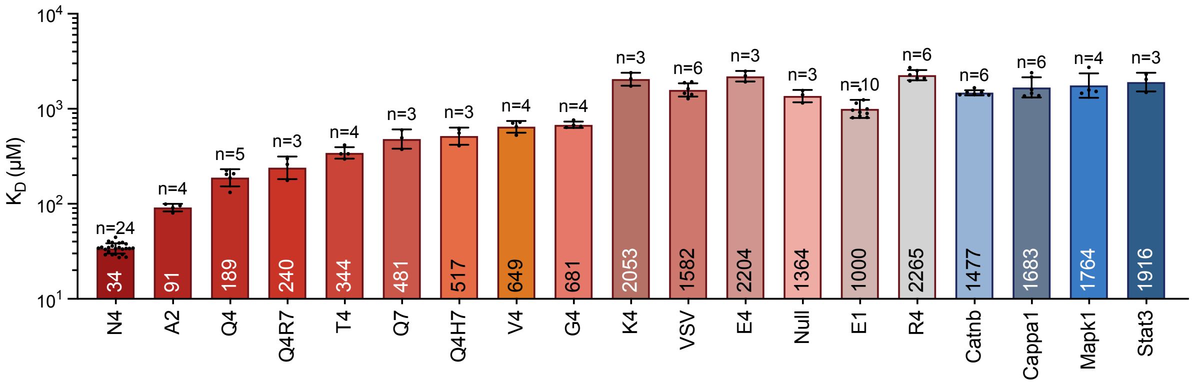


Figure S1: **Apparent $K_D$ values for OT-I specific peptides**. The $K_D$ values were obtained by fitting the steady-state binding using a 1:1 model with $B_{max}$ constrained to value obtained from the B2M or Y3 antibody binding using standard curve (see Fig. 1 and Methods). Geometric mean is displayed within the bars, mean values, error and N are listed in Table 1.

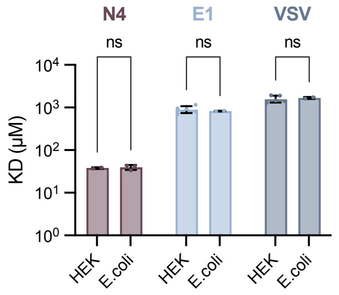


Figure S2: **The OT-I TCR displays similar affinity to pMHC produced in *E.coli* and HEK293T cells.** The *E.coli* produced pMHC where produced in-house whereas the HEK293T produced pMHC were supplied by the NIH tetramer facility.

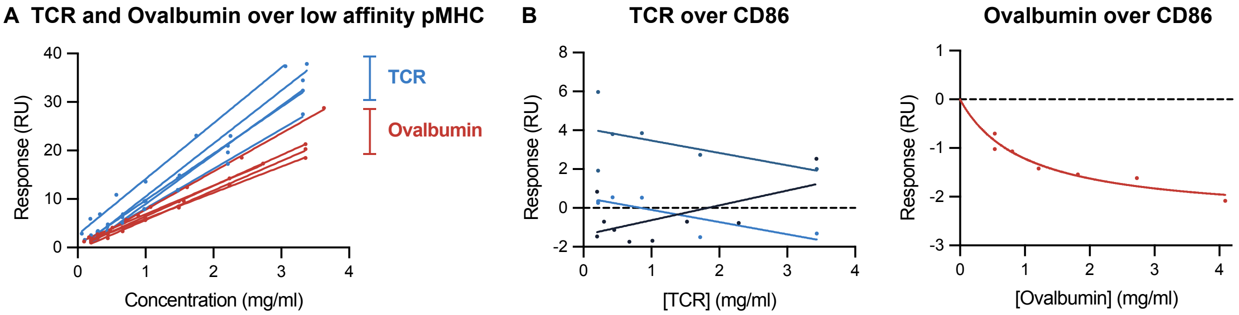


Figure S3: **An irrelevant protein (ovalbumin) can bind heterotrimic pMHC but not monomeric CD86. (A)** Steady-state binding of ovalbumin or OT-I analytes over pMHC surfaces. Note that the TCR displays higher binding than Ovalbumin suggesting that it binds both inactive and active pMHC whereas Ovalbumin binds only inactive pMHC. **(B)** Steady-state binding of TCR (left) or Ovalbumin (right) over CD86 surfaces reveals no detectable binding. Negative values can arise from modest refractive index effects.

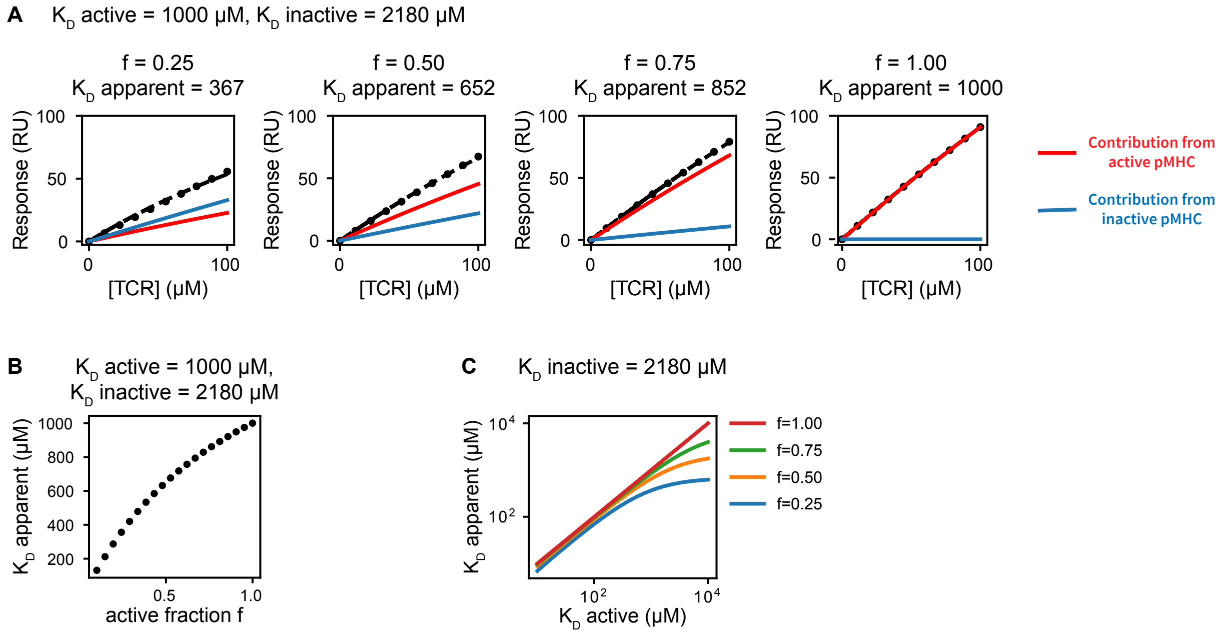


Figure S4: **Simulations highlight the impact of the fraction of active pMHC (f) on apparent $K_D$ values. (A)** Simulated TCR binding curves to surfaces containing variable amount of active and inactive pMHCs (columns). The overall binding signal (black) was fit to a 1:1 binding model to estimate the apparent $K_D$. **(B)** Apparent $K_D$ values derived from fitting the simulated curves in (A) plotted against the fraction of active pMHC (f). The apparent $K_D$ diverges from the active $K_D$ as f decreases. **(C)** Apparent $K_D$ values plotted against the true active $K_D$ for different f, demonstrating the dependency of apparent $K_D$ on both f and the active $K_D$.

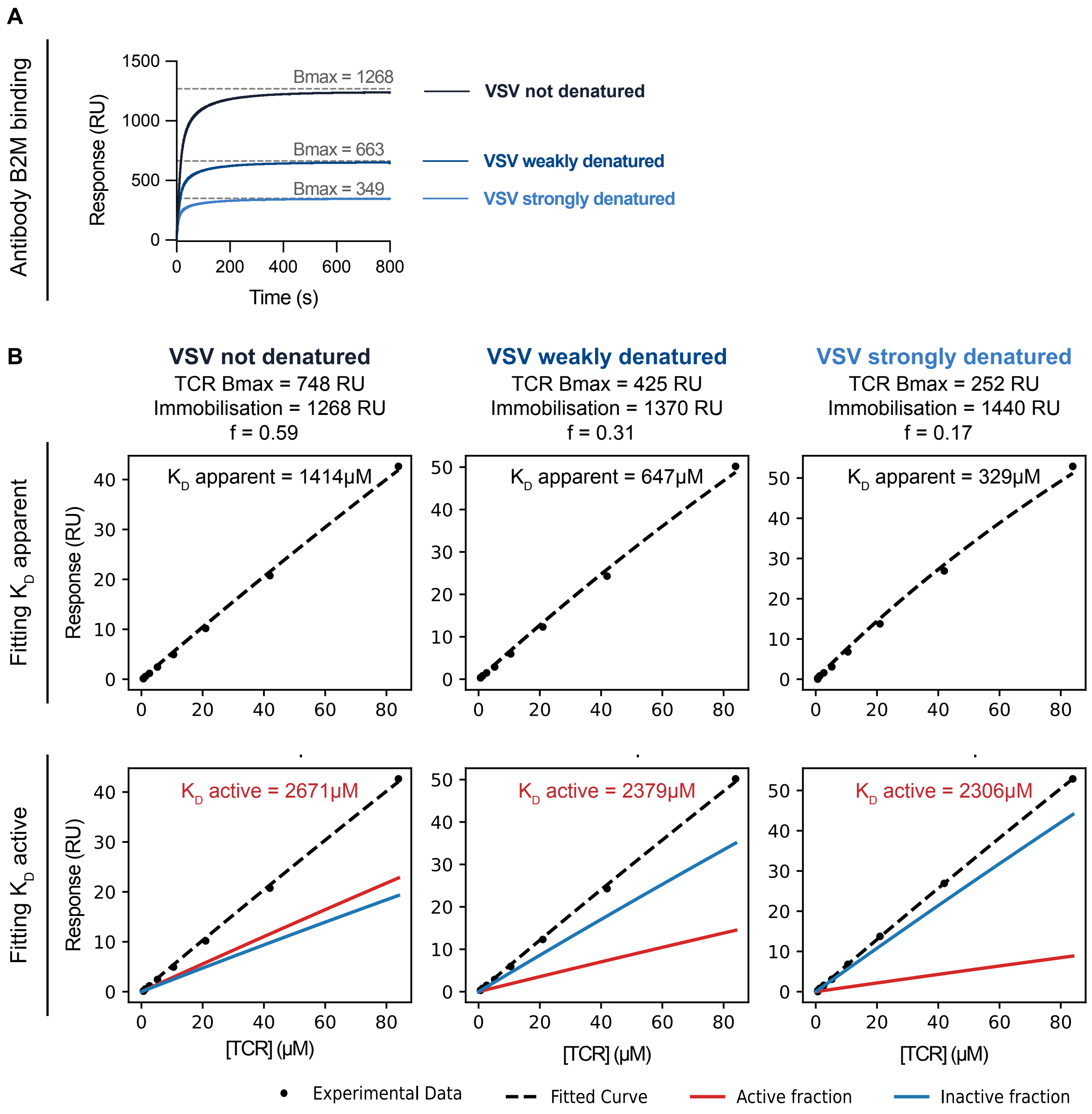


Figure S5: **Calculating the active $K_D$ provides similar results for OT-I binding to surfaces with large differences in the fraction of active pMHC.** The VSV pMHC was immobilised at similar levels on 3 flow cells in SPR before inducing denaturing by a short (weakly denatured) or long (strongly denatured) injection of glycine solution (pH 1.7). **A** B2M antibody binding curves to the three VSV pMHC surfaces. The TCR $B_{max}$ was estimated using the standard curve in Fig. 1B. **B** Steady-state TCR binding response to the 3 surfaces (columns). To determine the apparent $K_D$, the data was fit with a 1:1 binding model with constrained $B_{max}$ to determine $K_D$ apparent (top row). To determine the active $K_D$, the workflow in Fig. 2G-I was used, where the fraction of active pMHC was calculated from the ratio of TCR $B_{max}$ to pMHC immobilisation, and $K_D$ inactive was fixed to 2180 $\mu$M. While the apparent $K_D$ displayed large differences, the active $K_D$ produced consistent results across all VSV pMHC surfaces.

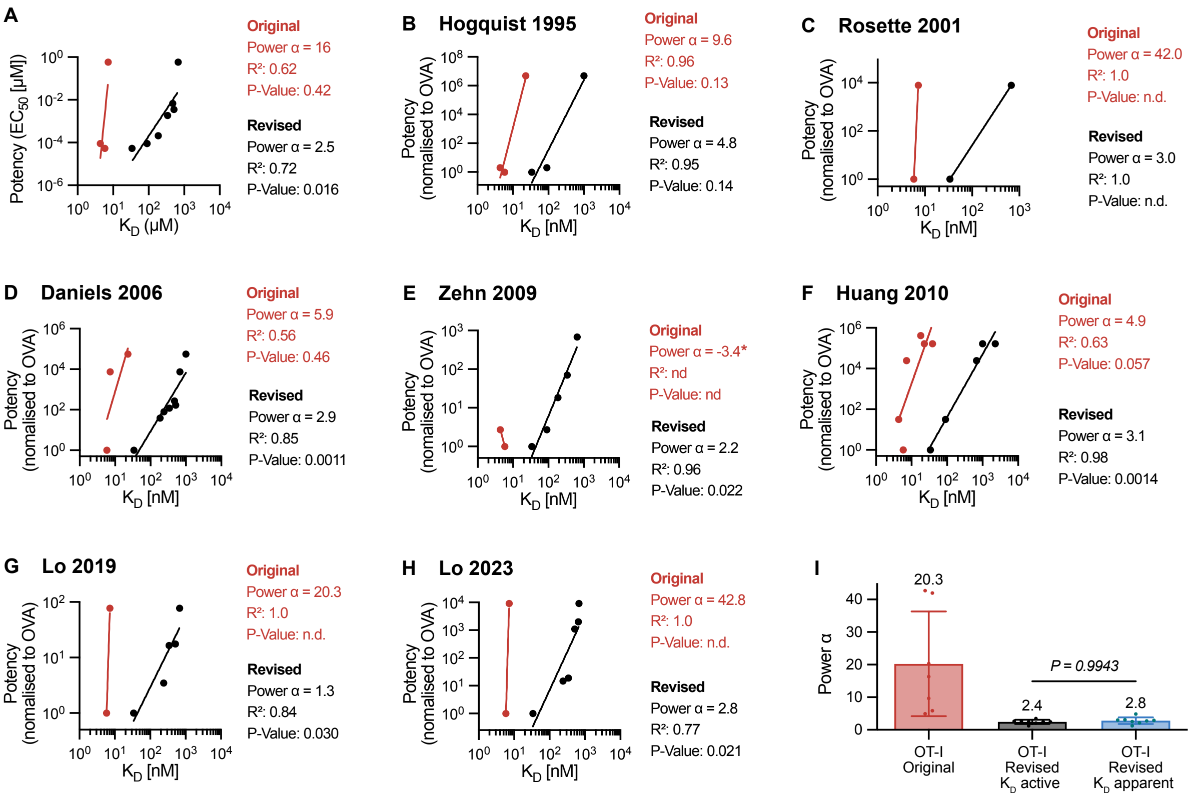


Figure S6: **Discriminatory power of OT-I TCR calculated with $K_D$ apparent shows imperfect discrimination.** Plots show peptide potency over original or apparent $K_D$ values. **(A)** Potency data from Fig 5A. Data is mean of N=2 independent experiments. **(B-H)** Published potency data from the indicated study over original or apparent $K_D$ values. A power law (potency $\sim (K_D)^{\alpha}$) is fit to the data to estimate the discriminatory power $\alpha$. A Pearson correlation is used to determine $R^2$ and p-values on log-transformed values. **(I)** The discriminatory power from panels A-H in comparison with discriminatory power calculated with active $K_D$ values. The p-value is determined using a t-test.

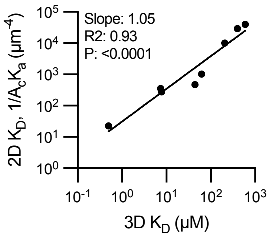


Figure S7: **Quantitative comparison of 2D and 3D affinties for the 1E6 TCR.** The log-transformed data was fitted with a linear regression. An F-test was used to determine a p-value for the null hypothesis that the slope is equal to zero. All data was taken from Cole et al (39).
